# Micro-probing enables high-resolution mapping of neuronal subpopulations using fMRI

**DOI:** 10.1101/709006

**Authors:** Joana Carvalho, Azzurra Invernizzi, Khazar Ahmadi, Michael B. Hoffmann, Remco J. Renken, Frans W. Cornelissen

## Abstract

The characterization of receptive field (RF) properties is fundamental to understanding the neural basis of sensory and cognitive behaviour. The combination of non-invasive imaging, such as fMRI, with biologically inspired neural modeling has enabled the estimation of population RFs directly in humans. However, current approaches require making numerous *a priori* assumptions, so these cannot reveal unpredicted properties, such as fragmented RFs or subpopulations. This is a critical limitation in studies on adaptation, pathology or reorganization. Here, we introduce micro-probing (MP), a technique for fine-grained and assumption free characterisation of subpopulation RFs. Without specific stimuli or adapted models, MP mapped the bilateral RFs characteristic of observers with a congenital pathway disorder. Moreover, in healthy observers, MP revealed voxels that capture the activity of multiple neuronal subpopulations. Thus, MP provides a versatile framework to visualize, analyze and model, without restrictions, the diverse RFs of cortical subpopulations in health and disease.

## Introduction

Over the past decade, our understanding of human brain function, organization and plasticity has increased tremendously. An essential contribution to this success has come from the ability to characterize the receptive field (RF) properties of neurons. The first electrophysiological measurements of those receptive field properties (in monkeys and cats) showed that the visual cortex is retinotopically organized and contains multiple maps representing the visual field (1–3). The development of non-invasive neuroimaging techniques, such as fMRI, opened a window to study brain activity directly in humans, albeit at a somewhat coarser scale. A subsequent boost to the field of visual neuroscience came from the development of biologically plausible computational models, which enable detailed characterization, also in humans, of the collective stimulus-referred RF of a population of neurons (4). Such detailed characterization is essential for linking brain function and behavior and understanding brain plasticity (for reviews, see e.g. (5–7)). In recent years, the approach has been extended towards neural-referred pRFs (8) and other perceptual domains, such as audition and numerosity (9, 10).

The conventional population RF (pRF) approach requires making *a priori* assumptions about the spatial, temporal and feature-selective properties of the pRF. This limits its ability to reveal unexpected pRF shapes, properties and subpopulations. In addition, it assesses the aggregate response across all neuronal subpopulations present within a voxel and thus primarily represents the most vigorously responding subpopulation. To advance our understanding of visual processing and cortical organization, approaches that can capture more fine-grained properties of distinctive subpopulations would be required. In particular, characterization of the shape of RFs may reveal its selectivity and specificity (11–16). An example of a model that results in a detailed characterization of the RF structure is the single unit receptive field (suRF). By modelling the neuronal activity with Gabor functions, suRF enables estimation of the size of average single-neuron RF (17).

A model with minimal *a priori* assumptions – enabling advanced pRF-mapping techniques – could be used to study visual pathologies, which are often characterized by highly atypical cortical pRF shapes. In such conditions, asymmetrical or even fragmented pRFs can arise that severely challenge both conventional retinotopic and contemporary pRF mapping techniques (18, 19). Such conditions could be an important application of advanced mapping techniques. While atypical and even unexpected pRFs may arise in deafferented visual cortex due to retinal or cortical lesions, very systematic deviations have been found in congenital visual pathway abnormalities (20). In albinism, for example, the visual cortex receives input from both hemifields, resulting in voxels with bilateral pRFs in opposing visual hemifields (21, 22). These pRFs are associated with an erroneous projection of the axons from the temporal retina to the contralateral hemisphere, which affects the central vertical portion of the visual field. Due to the predictability of the resulting pRF-abnormalities, i.e. their bilaterally split shape, albinism is ideal for validating the performance of new pRF-mapping techniques that have been optimised – with minimal *a priori* assumptions – to reveal highly atypical pRFs.

We therefore developed a technique for capturing the activity and properties of neuronal populations and subpopulations, which we present here. This approach efficiently samples the entire stimulus space, such as the visual field, with a “microprobe”: a 2D Gaussian with a small standard deviation. Regions of stimulus space that exhibit better model fits will be more heavily sampled. Like the conventional pRF approach, these microprobes sample the aggregate response of neuronal subpopulations, but they do so at a much higher resolution. Consequently, for each voxel, the MP generates a probe map representing the density and variance explained (VE) for all the probes. The probe maps are visual field coverage maps that can be used for visual inspection and for directly deriving neural properties such as symmetry. Moreover, following probe thresholding and clustering, they can also be used for identifying multi-unit receptive fields (muRFs). The muRFs properties can be further characterized by fitting shape models, if desired.

A primary advantage of this new approach is that it makes minimal *a priori* assumptions about the muRF properties or their number. For example, there is no need to specify up front the expected number of locations in stimulus space that a recording site (voxel) may respond to. We validated and tested the limits and capabilities of our new method using both in-vivo visual field mapping data and simulations. Without using specific stimuli or models, we recovered bilateral receptive fields in primary visual areas that are typical for the abnormal visual field representations in albinism (21, 23). Moreover, to demonstrate its versatility, we empirically estimated various muRF properties in healthy participants.

## Results

### Retinotopic mapping using MP

We applied MP to the retinotopic mapping data of healthy observers. Figure 1 shows three examples of probe maps for three representative V1 voxels and the derived muRF properties. It shows the estimated muRFs at various k-thresholds (which sets the percentage of probes with the strongest VE included in the muRF estimation) and compares these to the conventional pRF estimates.

**Figure 1.**
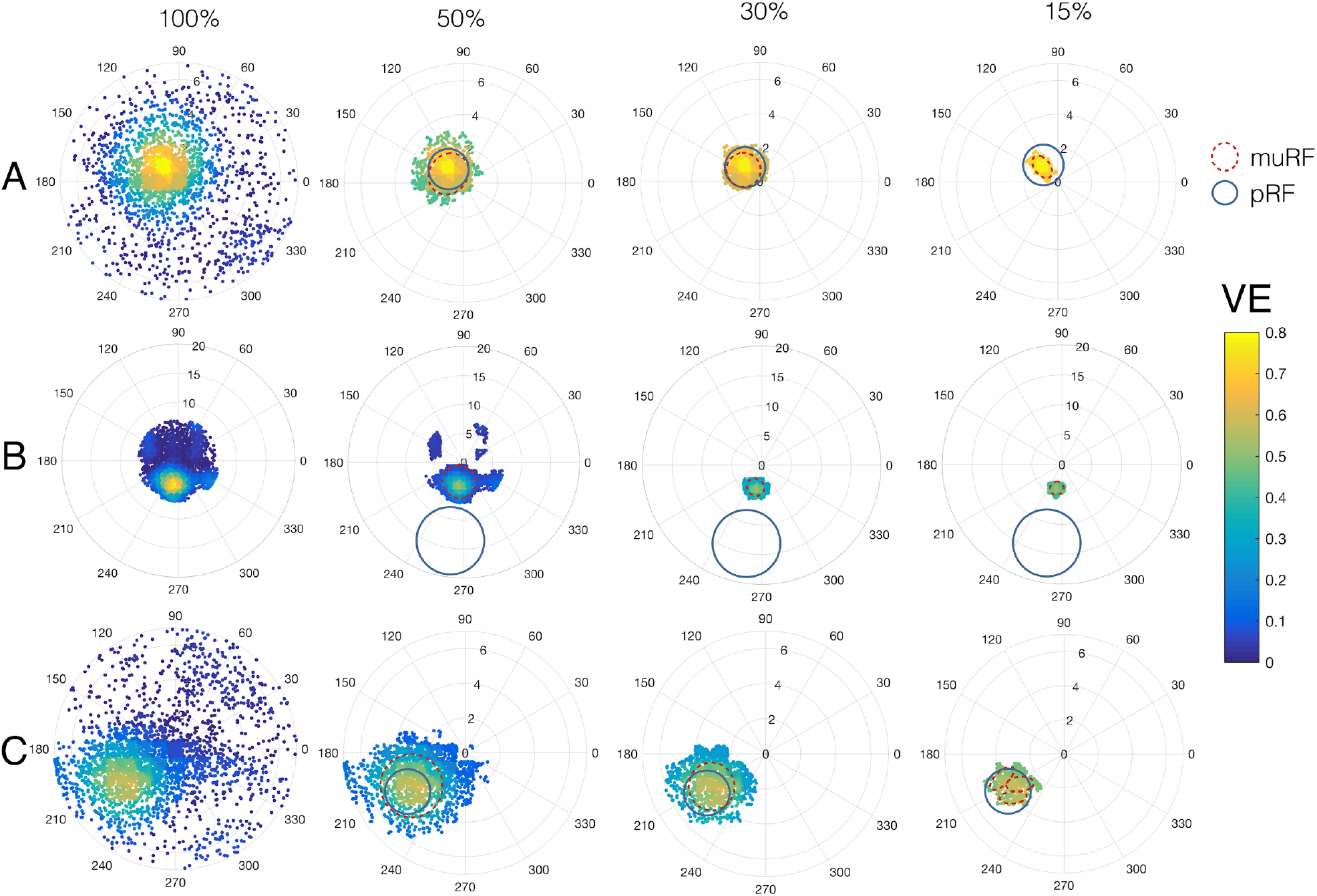
Examples of MP probe maps. Also shown are comparison of MP derived muRFs and conventional pRF estimations for V1 voxels. Shown are results obtained for three V1 voxels and with probe maps thresholded at a k-threshold of 100%, 50%, 30% and 15% (this k-threshold determines the percentage of probes with best VE included in the muRF estimation). It is clear that the conventional pRF (blue circles) and MP-based muRF estimates (dashed red outlines) can differ in various ways: A) estimated muRF size and shape depend on k-threshold. B) MP found a muRF with a high VE (0.49), while the estimated pRF had low VE (0.005) and was located outside of the stimulated region. C) At lenient k-thresholds (100%, 50%, 30%), MP revealed a single muRF, while at a more stringent threshold it detected multiple muRFs. Data was obtained during retinotopic mapping. The V1 voxels were extracted from the right hemisphere of observer S07.

Figure 1 indicates various features of the MP approach. First, it shows that the estimated size of the muRFs depends on the chosen k-threshold. More liberal thresholds result in muRFs that are approximately similar in size to conventional pRFs, whereas more stringent thresholds result in smaller muRFs (Figure 1A). Figure 1B shows an example in which MP resulted in accurate detection of the muRF, whereas the pRF estimate was not. Note that the location of the muRF does not depend on the chosen threshold. A more stringent threshold also enables identification of multiple muRFs (Figure 1C). Using simulations (Figure S1), we confirm that a stringent k-threshold minimizes the eccentricity error, while more lenient ones minimize the size error. Polar angle estimates were not influenced by the k-threshold. Additional simulations showed that: 1) the MP is highly robust to noise and can accurately determine the number of muRFs as well as their position and shape (Figure S2). The main factors affecting the accuracy of MP are the actual number of simulated muRFs and their proximity (Figure S3).

Figure 2A depicts the similarity of the eccentricity maps obtained with the pRF and MP approaches. Figure 2D shows that the muRFs and conventional pRF eccentricity are highly correlated. In the periphery, however, the estimated muRFs have somewhat lower eccentricity (i.e. they are situated more foveally) than the accompanying pRFs. This is particularly noticeable for higher-order areas.

**Figure 2.**
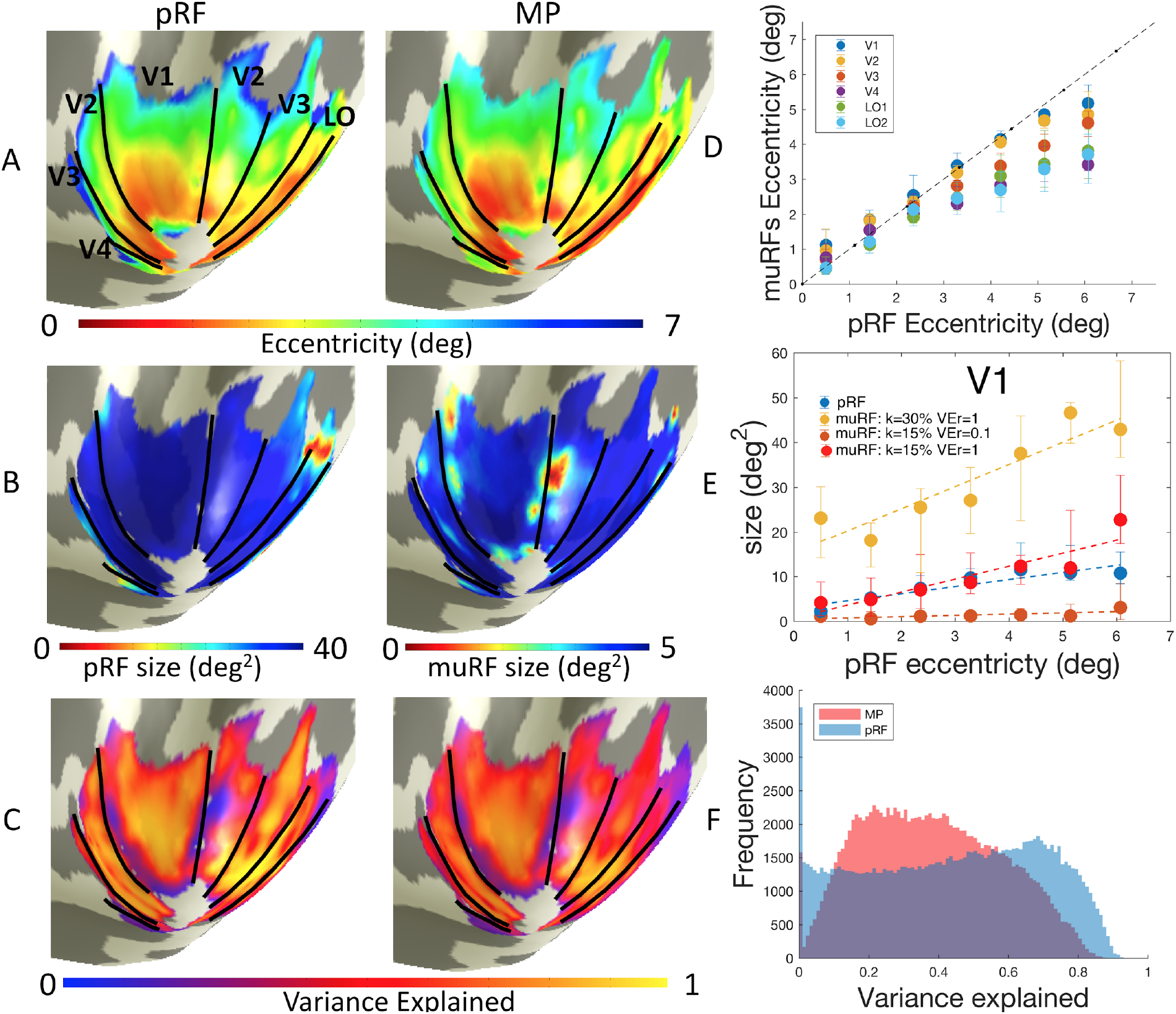
Comparison of MP-derived muRF and conventional pRF estimates. Panel A: pRF and muRF eccentricity maps projected on an inflated brain mesh. If MP identified multiple muRFs for a voxel, the eccentricity map shows the eccentricity of only one (arbitrarily chosen) muRF. Panel B. Left: pRF and muRF area maps projected on an inflated brain mesh. Panel C. Left: Comparison of MP-derived muRF and conventional pRF estimates VE. Panel D: Median eccentricity of the muRFs as a function of the eccentricity of the conventional pRF (the dashed line represents a perfect correlation). The pRF eccentricity was binned in 1 degree bins of eccentricity (data from 7 healthy observers: 14 hemispheres). Error bars represent 5% and 95% confidence intervals. For the visual areas tested, muRF eccentricity correlated highly with that of the corresponding conventional pRF (correlation coefficients vary between 0.98 and 1, depending on the visual areas, with p-values <0.05; see Table S1 for the correlation values and corresponding p-values). Figure S4 shows the relationship between the eccentricity of all muRFs and the pRF. Panel E: muRF (at different k-thresholds and VEr (VE-range)) and pRF size as function of eccentricity. The muRF size of an arbitrarily chosen muRF was binned in 1 degree bins of eccentricity (data from 7 healthy observers: 14 hemispheres). Error bars represent 5% and 95% confidence intervals. The dashed lines represent the linear fit. Figure S5 shows the relation between the size and eccentricity of all muRFs and the 6 visual areas tested. Panel F: Histogram of the VE for muRFs (blue) and pRFs (red). The VE was based on the cumulative activity of the number of muRFs. The histogram shows the data accumulated across 6 visual areas and 7 healthy observers (14 hemispheres).

Figure 2B shows the projections of pRF and muRF size on an inflated brain mesh. Due to our choice of k-threshold (15%) and a VEr (maximum difference in VE between the most and least explanatory probe) of 0.1, the muRF sizes shown here are significantly smaller than those of the pRFs (note the different scales). Nevertheless, Figure 2E shows that both pRF and muRF size increase with eccentricity, irrespective of the k-threshold used. Note how the choice of k-threshold influences the estimated muRF size. Figure S5 shows the same effects for a number of visual areas.

Figure 2C shows (projected on an inflated brain mesh) the close similarity of the VE for pRF and muRF model estimates. Figure 2F shows how MP performs better than the conventional pRF for voxels with low explanatory power (VE<0.1). In contrast, for voxels with a very high explanatory power (VE>0.8), the conventional pRF has a higher VE than MP. This is partly because the conventional pRF method tends to estimate larger pRFs, which also results in a higher VE. Figure S6 shows that increasing the k-threshold for MP also increases the estimated muRF size, resulting in a higher VE.

### Application of MP in albinism

To demonstrate the biological relevance of our new technique, we applied MP to data obtained in observers with albinism. Based on previous work in observers with albinism, we expected to find mirror symmetry in the positions of the estimated muRF with respect to the vertical meridian (21–23). Figure 3A shows the projection of the symmetry coefficients (calculated based on the probe maps regarding the vertical midline) onto the reconstructed hemispheres of a representative observer with albinism and a control observer. See method section “Symmetry analysis of probe maps” for more details. In albinism, the probe maps revealed a large number of voxels with muRFs that were mirrored across the vertical meridian. Closer inspection of the probe map of Figure 3C, showed highly symmetrical and spatially organized muRFs in an example voxel. The cortical projections showed that most of the symmetry coefficients were much higher in albinism than in the control observer, and that neighbouring voxels had similar symmetry coefficients. Central regions showed higher symmetry coefficients than peripheral ones. Moreover, we found that a clear overlap between the cortical region with high symmetry values and the right-left hemifield overlap cortical region (dashed line) that was determined based on stimulating the left and right hemifield in separate experiments (described in (24)). In control observers, high symmetry values were found for voxels with a muRF near the border of visual areas (e.g. V1/V2), where the muRF is expected to be located on or very close to the vertical meridian. Figure 3B illustrates why such voxels also have high symmetry coefficients.

**Figure 3.**
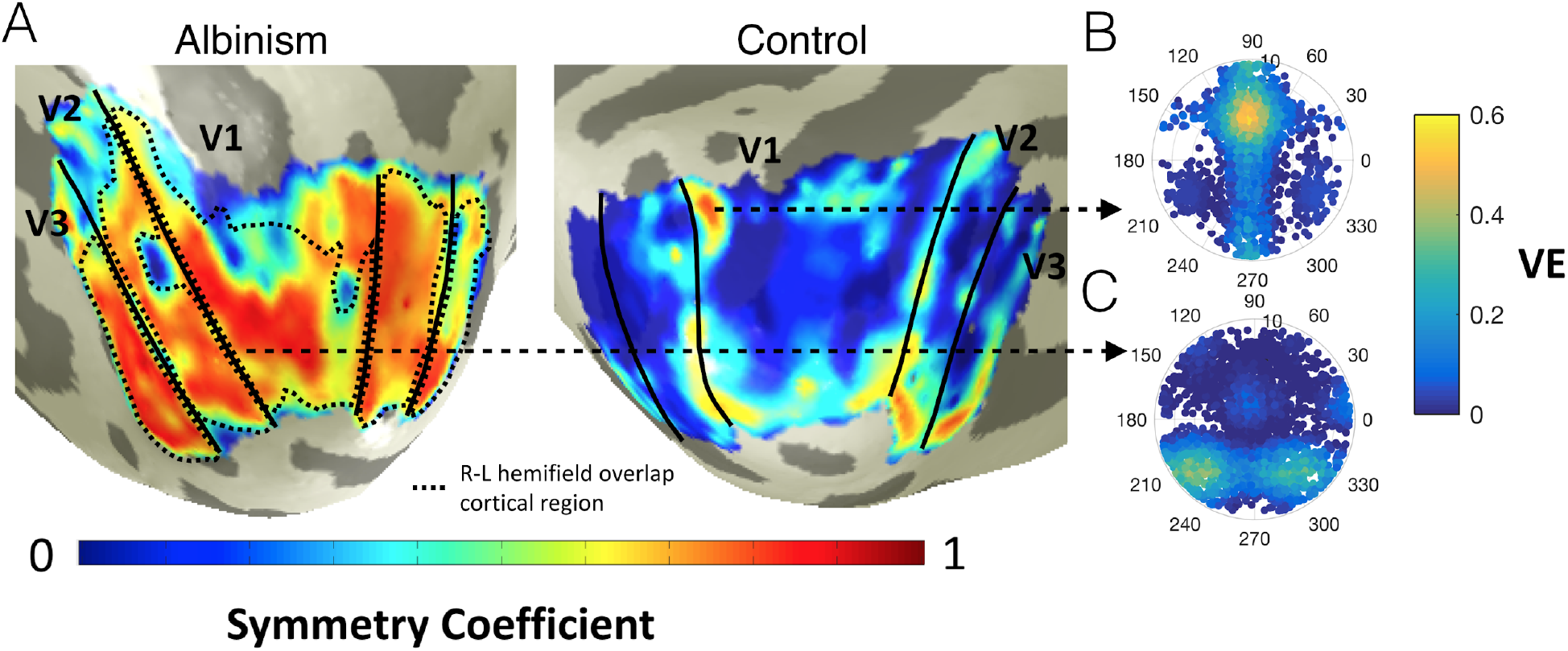
Symmetry maps in albinism and aged-matched control. Symmetry map for the left hemisphere of the observer with albinism A03 and of the aged match control C03. The black continuous lines outline the visual areas and the black dashed line outlines the misrouted cortical region calculated based on the overlap of right and left hemifields (Hoffman et al, 2003; 2012). On the right are two example probe maps (k-threshold = 100%) for voxels of the control (top) and an observer with albinism (bottom). The thresholded probe maps at 100%, 50%, 30% and 15% and the estimated muRF are shown in Figure S7. Figure S8 shows that MP tends to perform better than the conventional bilateral pRF model for voxels with very low VE (<0.1), which is in accordance with the results for healthy observers.

To demonstrate the versatility of MP, Figure 4A shows the symmetry calculated for a series of symmetry axes for the V1 region of the right hemisphere of every observer during full field stimulation. Controls had slightly increased symmetry coefficients for both the horizontal and vertical symmetry axes (0 and 90 degrees). This reflects the symmetry of neuronal populations located along the vertical and horizontal meridians and the distribution of muRFs in the visual field. Figure S9 shows the high number of muRFs located on the horizontal meridian. For observers with albinism, the inter-observer variability corresponded with their differing levels of misrouting. As expected, those with severe misrouting (top row) showed a high degree of symmetry for the vertical axis. No systematic differences were found for albinism observers with low levels of misrouting (bottom row). The V1 symmetry coefficients for the vertical axis correlate highly with the clinically established level of misrouting (Figure 4B). The symmetry coefficient to the vertical meridian is thus indicative of misrouting.

**Figure 4.**
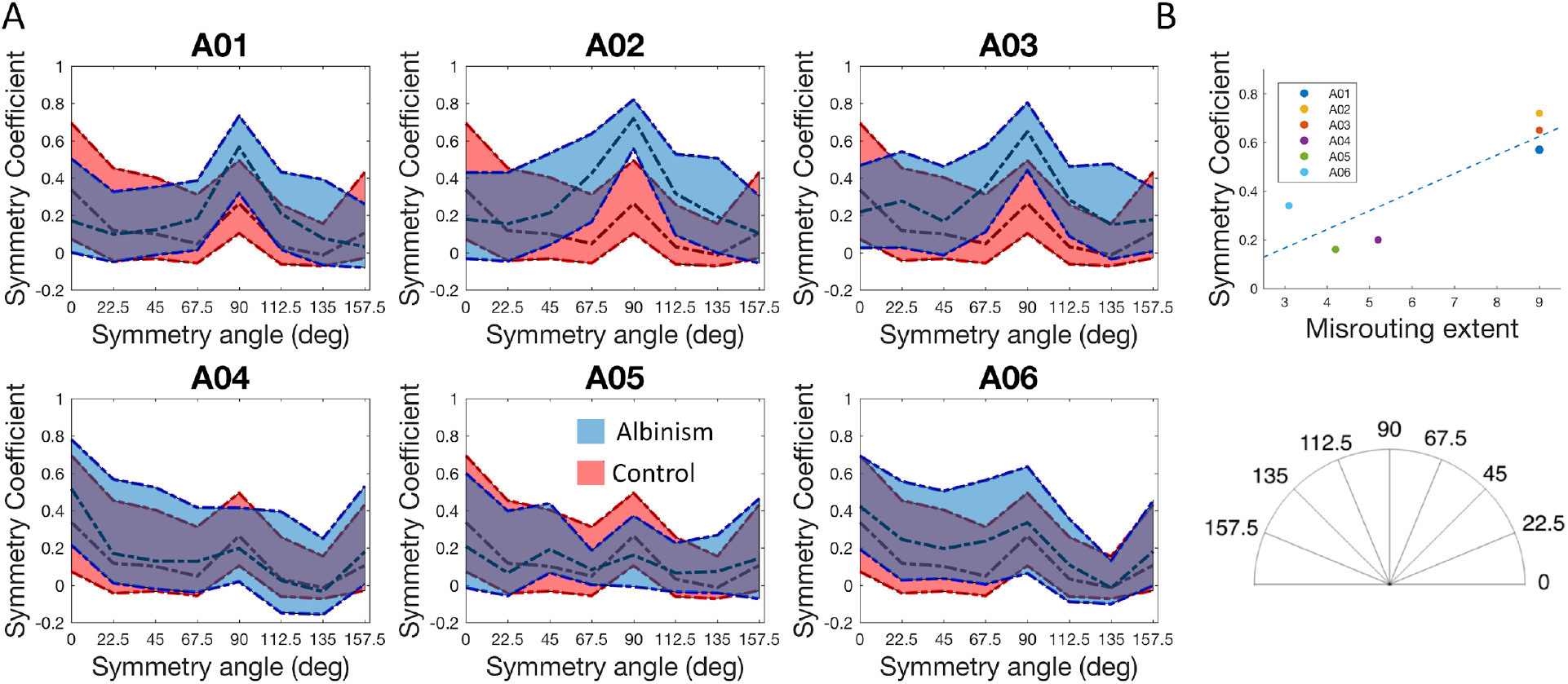
Symmetry coefficients based on probe maps. Coefficients in healthy controls and albinism are shown as a function of the angle of the symmetry axis. Symmetry coefficients for the V1 of the right hemisphere during full field stimulation calculated across 8 symmetry axes (see right inset). The dashed lines represent the 5%, 50% and 95% confidence intervals. Albinism distributions are shown in blue and controls in red. Given that the inter- and intra-observer variability in healthy controls was low (see Figure S9A), their symmetry coefficients were averaged. The albinism observers are shown in order of decreasing level of misrouting (see Table S2) as assessed independently (24). Panel B: Correlation between the symmetry coefficient to the vertical meridian and the mean misrouting extent of the observers with albinism based on independent stimulation of the left and right hemisphere (Table S2). The symmetry coefficients in the vertical meridian correlate highly with the mean misrouting extent calculated for the three visual areas analysed (V1 R=0.88, p=0.02; V2 R=0.77, p=0.07; V3 R=0.84, p=0.04).

### Using MP to estimate muRF properties

Using MP, it is relatively straightforward to explore a variety of muRF properties, such as the number of muRFs per voxel, muRF bilaterality or muRF shape (e.g. their elongation). Figure 5A shows a map of the spatial organization of the number of muRFs over the visual cortex, projected on the inflated right hemisphere of a representative observer. Neighbouring voxels tend to have a similar number of muRFs. Comparable results were also observed in observers with albinism (Figure S10A). Closer inspection of the probe map of a single voxel (5B) shows how MP resolves multiple muRFs and their corresponding properties. Figure 5C shows how muRFs tend to be more elongated (i.e. less spherical) in the (para-)fovea compared to the periphery. We observed this trend in all visual areas analysed (Figure S11A). Figure 5D shows how unilateral and bilateral muRFs were distributed over the visual cortex, again for a representative observer. Closer inspection of the probe map of one voxel (figure 5E), shows how MP revealed two muRFs situated in opposite hemifields and quadrants. For the vast majority of the voxels, the estimated muRFs were located within the same (contralateral) hemifield (dark blue). However, some voxels contained bilateral muRFs. These are muRFs that process information from both the left and right hemifields. The bilateral muRFs appeared to be spatial organized and clustered along the vertical meridian (red blobs in Panel D). Panel F shows the histograms of the bilateral muRFs, which peak near the vertical meridians, conforming this observation.

**Figure 5.**
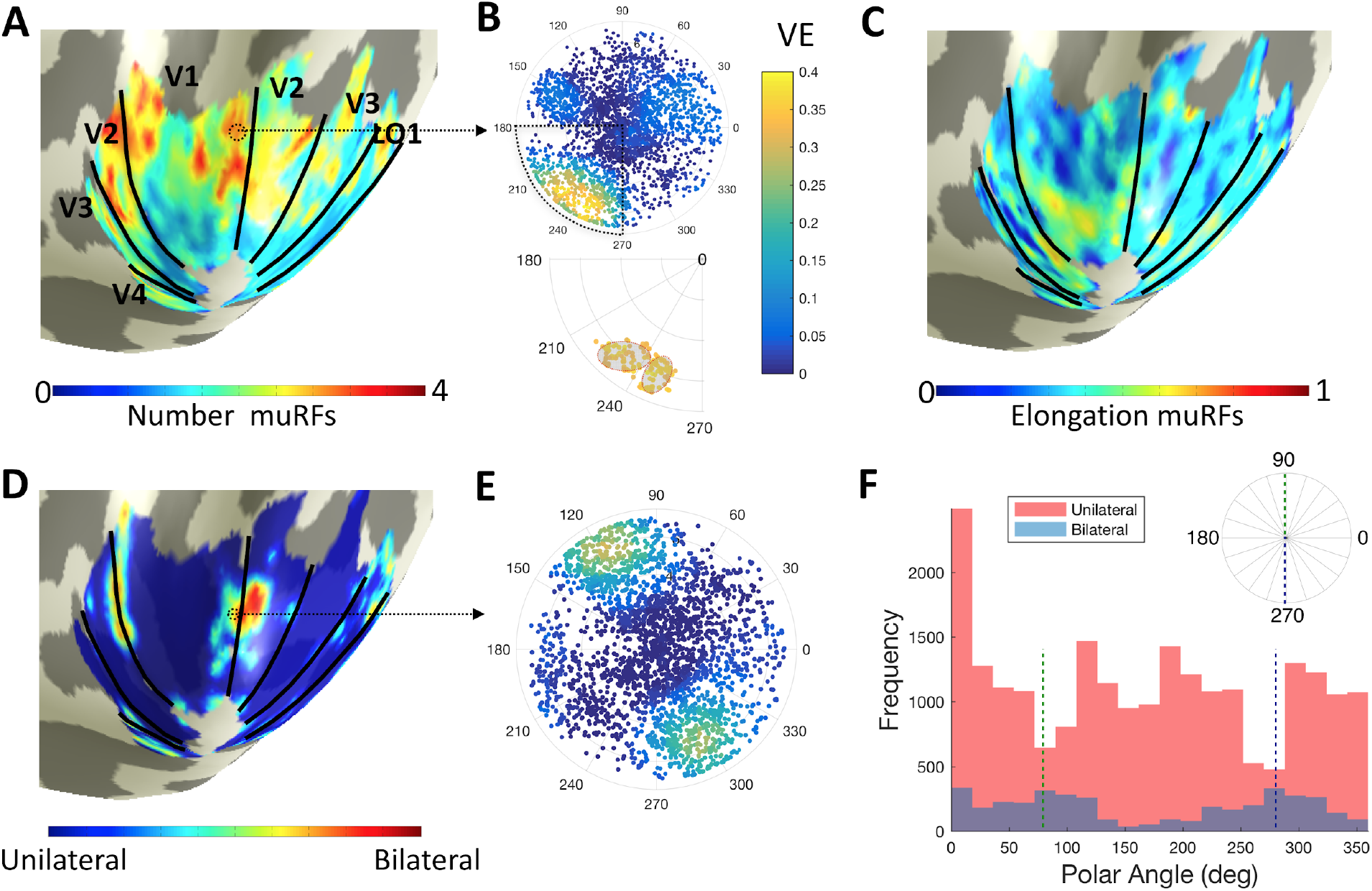
Examples of muRF properties estimated in the visual cortex of healthy observers. Panel A: Projection on an inflated brain mesh of the number of muRFs estimated per voxel (right hemisphere of observer S07). Black lines represent the borders of visual areas. Panel B, upper part, shows an example of a MP map of a V1 voxel (location indicated by the dashed circle in Panel A; the lower part shows a zoomed-in view of one quarter field of the probe map indicating the estimated muRFs (outlined by a red dashed line). Panel C: Projection on an inflated brain mesh of the muRF elongation (right hemisphere of observer S07). Black lines indicate the borders of visual areas. Panel D: Projection on an inflated brain mesh of the unilateral-bilateral label, estimated per voxel. The intermediate colors between blue and red resulted from interpolation during mesh projection. Panel E: Probe map of a representative voxel with a bilateral RF (this was not mirrored in the vertical meridian, which differs from observers with albinism; location is indicated by the dashed circle in Panel D). Panel F: Histogram of the number of unilateral (red) and bilateral (blue) muRFs in V1 (14 hemispheres, 7 healthy observers). The dashed line depicts the vertical meridian. The regularity of the shape can be assessed by measuring the skewness of the distributions. V1 has predominantly regular shapes (skewness = 0). The irregularity of the shape tended to increase with visual hierarchy (Figures S11B and D). The kurtosis of the probe distribution was positive. This indicated that there was a high sample density at the pRF center and that the thresholding was not too severe (Figures S11C and E).

## Discussion

In this study we introduced microprobing (MP), a versatile, model-based fMRI analysis framework that requires only minimal *a priori* assumptions about the underlying biological mechanisms. By repetitively applying microprobes to the fMRI time series, we produced a probe map that reveals the detailed visual field coverage of each voxel. These maps enable the extraction of fundamental muRF-properties that were previously not accessible, outside single cell recordings. We validated our approach using both empirical and simulated data and demonstrated its biological validity by revealing the highly atypical visual field representations for observers with albinism, without making any prior assumptions about this. Finally, we demonstrated how various receptive field properties, such as shape and position, can be determined with relative ease for the entire visual cortex.

### MP recovers multiple RFs within a single voxel

Our analysis of empirical data of observers with albinism and simulations both revealed that MP accurately detects – for each voxel – the number of muRFs as well as their position and shape. Applied to observers with albinism, MP resolved muRFs mirrored in the vertical meridian, thus revealing the simultaneous processing in one hemisphere of the signals coming from both the contralateral and ipsilateral hemifields. This corroborates previous studies that took the bilateral representation of the RFs as an *a priori* starting point (20, 21, 23). Importantly, MP does not require making such an assumption. Instead, based on stimulation across the entire visual field, it quantifies the degree of symmetry in the vertical meridian (or other directions) in the probe maps.

As the next step, we used the symmetry values to quantify the extent of misrouting and identify the misrouted cortical region per observer. Remarkably, in controls, we showed that highly symmetric probe maps delineate the borders of the visual areas. The fact that MP revealed the atypical visual field representations in albinism suggests that these muRFs found in controls are also biologically genuine and meaningful. MP therefore enables a straightforward estimation of atypical RF representations without requiring additional stimuli or assumptions.

In healthy observers, we demonstrated that MP not only resolves multiple muRFs within a voxel, but also their properties. The muRFs are spatially organized and the number of muRFs increases with eccentricity. Note that due to cortical magnification, it is likely that the subpopulations are more widely spread in the periphery than in the fovea, and thus easier to identify. We found that approximately 10% of the estimated muRFs were bilateral and located near the vertical meridian. This supports the hypothesis that these bilateral muRF may derive from visual callosal connections that contribute to the integration of the cortical representation at the vertical midline (25–28). Previous studies focusing on the medial superior temporal (MST) area also reported bilateral RFs (29–31). We also found multiple mirrored muRFs (> 2) in albinism, corroborating the finding in healthy observers (Figure S10). This suggests two possible explanations that require further study: 1) that neurons may simultaneously process information from distinct portions of the visual field and 2) subpopulations with spatially distinct properties may be present within a single voxel.

### MP is robust and recovers biologically meaningful RF properties with only minimal prior assumptions

The probe maps revealed that the muRFs can be heterogeneous in shape. MP enables muRF shape estimation without assuming specific shape properties *a priori*, such as circular symmetry. By assessing the statistical properties of the probe distributions, we showed how muRF shape can be characterized. Based on such assessments, we found that the majority of the muRFs tend to be elongated. This is in line with recent studies that found that the pRFs tend to be elliptical and radially oriented towards the fovea (15, 16). These findings support the functional differentiation of visual processing from fovea to periphery and across the ventral and dorsal cortical visual pathways.

By comparing the characteristics of muRFs to those of the conventional pRF, in healthy observers, the following three conclusions can be drawn. Firstly, the eccentricity estimates for muRFs and pRFs correlate closely. Our proposition that muRFs are biologically meaningful is supported by the similarity between the eccentricity maps obtained with pRF and MP, the fact that muRF size increases with its eccentricity. In the periphery, however, eccentricity estimated by muRF tends to be smaller than eccentricity estimated by conventional pRF. This could be explained in part by the fact that muRF size also tends to be smaller, which corroborates previous work showing that the smaller pRFs estimated for an orientation contrast stimulus also result in lower eccentricity estimates for the same voxels, especially in higher-order areas such as lateral occipital cortex (32). It may also reflect different model specifications. By design, the MP estimates are located within the stimulated visual field. Consequently, the muRFs estimates based on the probe maps are also situated within the stimulated part of the visual field. In contrast, the conventional pRF model allows for partially stimulated pRFs, the centers of which may be located far outside the stimulated visual field.

Secondly, the level of specificity with which muRFs are estimated is defined by varying the k-threshold and VE-range in the MP. This ranges from many multi-units (very restrictive k-threshold and VE-range) to more aggregate responses (more lenient k-threshold and VE-range). At the fairly restrictive k-threshold of 0.15 and VE-range of 0.1, the MP approach estimated smaller muRFs than the conventional pRF. Such relatively small RFs are in agreement with other studies that also estimated significantly smaller pRF sizes than the conventional approach, for example using model-free approaches such back projection (15, 17, 33). Thirdly, compared to conventional methods, MP improves capture of the dynamics of the measured signal, especially for voxels with signals that have a low explanatory power. We also determined that the muRF estimations are robust to noise, as MP could be performed reliably despite the presence of nystagmus in the observers with albinism. This robustness to noise was also confirmed using simulations.

### Limitations

We identified multiple muRFs in voxels in healthy observers, but the functional implications of this finding are uncertain. We have assumed that muRFs have biological relevance, but some of the units may have resulted from artefacts related to segmentation, from voxels stranded on the cortical sulci, or from partial voluming. There are several ways in which the effect of such artefacts could be reduced: correcting for partial voluming (34), identifying and extracting local sulci (35, 36) and applying MP to higher resolution functional data (< 1mm isotropic). Using more precisely controlled stimuli will contribute to unravelling the biological significance of muRFs and characterizing the neuronal subpopulations specialized in the processing of specific spatial and temporal properties (orientation, spatial frequency, colour etc) (32, 37, 38).

At present, MP is a computationally intensive approach when compared to the conventional pRF model. We expect that software optimization and advances in hardware will contribute to reducing the computation time. We currently address this issue by using parallel GPU computing. Furthermore, the use of MCMC sampling is needed only to speed up the process/ limiting computing resource use, but is not fundamental to MP. In principle, probe maps could result from systematically probing every position in stimulus space, creating a densely covered probe map for each voxel. The use of a Markov-Chain means that the current probe maps contain more probes for regions with higher VE. Our current estimate of muRF shape is based on the clustering of the probes weighted by their VE. In contrast, pRFs are fitted to explain the VE of the signal. This explains why the muRFs shapes are sometimes different (e.g. more elongated) from the pRFs. In our view, neither is necessarily correct; they are just different ways to assess shape. Future work will be required to indicate which approach best approximates biological reality.

### Future directions

MP as presented here is a reliable and versatile method to study cortical organization, but it can be improved in several ways. First, using more efficient stimulus designs, such as a narrower bar or multi-focal stimulation (39, 40), could improve the performance of MP. Secondly, applying additional advanced data-driven metrics to extract the shape and number of muRFs may result in a more detailed characterization of the RF. Thirdly, the definition of a probe could be extended, for example to a difference of Gaussians, which may enable MP to also account for negative blood oxygen level dependent (BOLD) activity (41).

Previous studies have shown that pRF properties are not stable, and may change in response to environmental factors (stimulus, task) and cognitive factors such as attention (32, 37, 38). Given that the probe maps reflect the scatter in the location of the receptive field centers, this scatter may reveal the dynamic properties of the muRFs (changes in position) and could be related to the connections underlying the RFs. Moreover, in principle, MP could be extended to cortico-cortical models, such as connective field modeling (8), which would also enable high resolution mapping of the flow of information between brain areas.

Changes in pRFs have been reported in health and disease (for a summary see (7)). Regarding disease, this concerns differentiation following cortical and retinal lesions, schizophrenia and autism spectrum disorder. We anticipate that the application of MP to ophthalmologic and neurologic disorders as well as to adaptation studies will reveal additional characteristics of the RF structure, such as number and shape.

The recent development of ultra-high field fMRI enables the in-vivo examination of the human brain at a mesoscale and can reveal previously unmapped columnar organizations. However, there is a need for methods that can extract more detailed information on the structure and function of the cortex from this high-resolution data. The application of MP to high-resolution functional data has the potential to reveal how the muRFs and their properties are distributed across cortical depth. This will be crucial to study the functional differentiation of the visual processing across laminae. Moreover, it may complement previous studies of cortical organization across cortical depth, ocular dominance and columnar pinwheel organization for orientation selectivity (42–44).

In this study, we described the application of MP to characterize the spatial organization of the visual cortex. However, MP could also be applied to visual feature dimensions, other sensory modalities and cognitive spatially organized features such as numerosity (9, 10).

## Materials and methods

The methods are presented in the following order. First, we will go through the steps of the MP framework. Second, we will describe the acquisition procedure. Third, we will describe how the MP analysis is applied to simulations, to empirically acquired fMRI data from healthy observers, and to the fMRI data of a cohort of observers with albinism and age-matched controls.

### Micro-probing framework

MP uses Markov chain Monte Carlo (MCMC) sampling (see (45)) to efficiently sample the entire stimulus space. To describe the MP framework, we first define the probe with its variables and the corresponding priors used. Then we describe the muRF estimation steps. The framework is shown in Figure 6.

**Figure 6.**
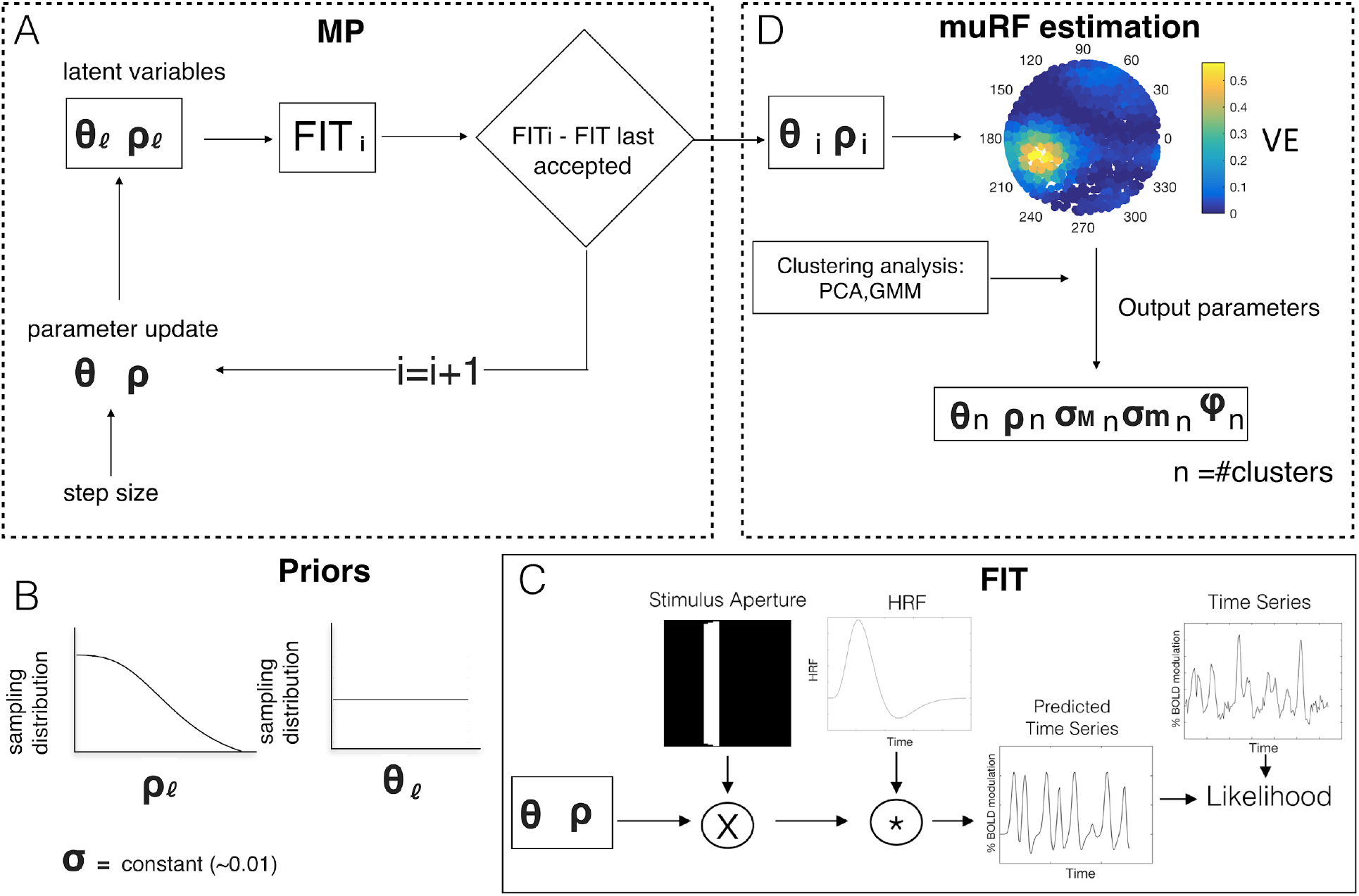
Overview of the microprobing framework. MP: creation of a probe map. FIT: Probe fitting procedure based on the conventional pRF approach (Dumoulin and Wandell, 2008). muRF: muRF estimation based on the probe map. Priors: *a priori* biological information about the distribution of the probes across the visual field according to *ρ*,*θ* and *σ*.

### Probe, latent variables and priors

Each probe was defined as a 2D Gaussian in Cartesian coordinates (in deg) in visual space, centered at 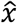 and 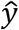, with a narrow fixed width (*σ*). The results shown in this study were obtained with *σ* = *0*.*01* deg.

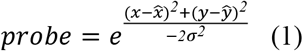

The center of the probe was defined using two latent variables. We used the nomenclature of Zeidman and colleagues (46). Let *l*_*ρ*_, *l*_*θ*_ be the latent variables corresponding to the radius and angle of the pRF center, respectively. The probe position of a RF in polar coordinates is given by:

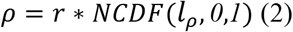

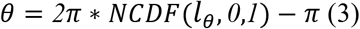

Here *r* is the radius of the stimulated visual field, in degrees. *NCDF* is the normal cumulative density function. In this study we initialized *l*_*ρ*_, *l*_*θ*_ with 0.5 and 1, respectively. Note that the centre of the probe is constrained to fall within the stimulated visual field.

To incorporate biological prior knowledge about the expected distribution of the probes within the visual field, a prior was assigned to each of the latent variables, *l*_*θ*_ and *l*_*ρ*_. Based on the work of Zeidman and colleagues (46), these priors were defined as normal distributions, *N(0,1)*. After conversion into polar coordinates (*ρ* and *θ*, equations 2 and 3), these priors express the assumption that the density of neurons is higher in the fovea than in the periphery (47), see Figure 6B. As the MP fitting procedure was done in Cartesian coordinates, polar coordinates were converted as follows:

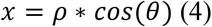

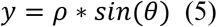

In the iterative MCMC procedure (Figure 1A), the next position (*i* + 1) was pseudo-randomly selected. The step size between the two probes was controlled by *d*_*proposal*_. In this study we defined *μ*_*d*_ and *σ*_*d*_ as 0.5 and 2, respectively.

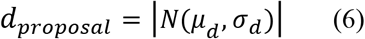

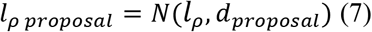

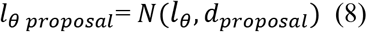

### Microprobe fitting

At each iteration of the MCMC, a probe position (defined by *ρ* and *θ*) was fitted. The fitting procedure resembles that of conventional pRF modeling (4); see Figure 6C. First, we predicted the voxel’s response to the stimulus *p*(*t*) by calculating the overlap between the stimulus and the probe at each time point *s*(*x*, *y*, *t*):

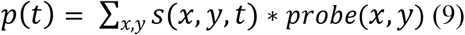

Second, we accounted for the delay in the hemodynamic response by convolving *p(t)* with the hemodynamic response function (48, 49). Finally, assuming a linear model of the fMRI response, we calculated the error per time point, *e*_*t*_, between the measured fMRI signal and the predicted signal using ordinary least squares fit.

Next, we calculated the likelihood, *l*_*t*_, associated with *e*_*t*_. Here we assumed that *e*_*t*_ is normally distributed, enabling the estimation of the mean and standard deviation (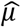 and 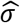, respectively). Given 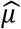 and 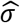, we calculated the total likelihood, *L*_*t*_, accounting for the contribution of the priors of *ρ* and *θ*.

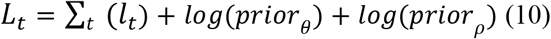

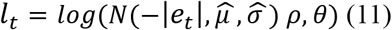

Finally, the likelihood of the current iteration was compared to the last accepted iteration, according to the following steps. First, the acceptance ratio, *Ar*, was computed.

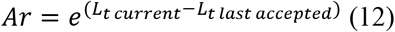

Second, a probability of random acceptance, *accept* was defined as N(0,1). Third, the Ar value was compared to the *accept* value: if the Ar was higher, the latent variables, *l*_*ρ*_, *l*_*θ*_, were updated. Based on the new *l*_*ρ*_, *l*_*θ*_ a new probe was defined and a new iteration took place.

To ensure that the entire visual field was probed, 12 different starting positions (equally distributed over the search grid) were defined. A total of 10000 iterations (∼ 833 per starting position) took place. This minimized the risk of local minima, i.e oversampling a specific muRF.

### Estimation of the muRFs

Following the iterations, we generated a probe map consisting of the projection in the stimulus space of all the probes weighted by their VE. Note that visual inspection of this probe map is informative in itself, regarding the properties of neural population that makes up the voxel. Based on this probe map, we can estimate various muRFs properties, such as their number, position, size, elongation, orientation, and irregularity of the shape and VE (Figure 6D).

The muRF estimation comprises three steps: first, we select the k% probes with the strongest VE (k-threshold). Additionally, the difference in VE between the most explanatory and the current probe must be lower than a given threshold (VEr). We found this improves any subsequent clustering and shape estimation. Unless otherwise specified, in the present study we used a value of 15% and 0.1 for k and VEr, respectively.

Second, the number of muRFs was determined by applying a weighted cluster analysis. Gap statistics were used to evaluate whether a single or multiple clusters were present (Tibshirani et al, 2000). In the latter case, to estimate the number of clusters, the Davies-Bouldin index clustering algorithm was applied (50). The maximum number of clusters that can be estimated needs to be defined a priori. In this study, we defined a maximum of four clusters for simulations and healthy observers and eight in case of observers with albinism and their aged-matched controls.

Third, the properties of the individual muRFs were determined using a Gaussian mixture model. This probabilistic model assumes that all data points were generated from a mixture of a finite number of Gaussians distributions of unknown parameters. The number of Gaussians to fit corresponds to the number of muRFs calculated in step 2. Furthermore, this model enables detection of the presence of a subpopulation within an overall population without requiring *a priori* identification of the subpopulation to which an individual probe belongs.

The code for MP is available on the visual neuroscience website https://www.visualneuroscience.nl/tools/.

## Empirical studies

### Participants and ethics statement

We recruited 7 participants (3 females; average age: 28; age-range: 26–32 years-old) with normal or corrected to normal vision. Prior to scanning, participants signed an informed consent form. Our study was approved by the University Medical Center of Groningen, Medical Ethical Committee and conducted in accordance with the Declaration of Helsinki.

### Data acquisition

Stimuli were presented on an MR compatible display screen (BOLDscreen 24 LCD; Cambridge Research Systems, Cambridge, UK). The screen was located at the head end of the MRI scanner. Participants viewed the screen through a tilted mirror attached to the head coil. Distance from the observer’s eyes to the display (measured through the mirror) was 120 cm. Screen size was 22×14 degrees. The maximum stimulus radius was 7° of visual angle. Visual stimuli were created using MATLAB and the Psychtoolbox (Brainard, 1997; Pelli, 1997).

### Experimental procedure

Each participant participated in one (f)MRI session. Retinotopic mapping was done using a standard drifting bar aperture defined by high-contrast contrast-inverting texture (4). The bar aperture moved in 8 different directions (four bar orientations: horizontal, vertical and the two diagonal orientations, with two opposite drift directions for each orientation). A single retinotopic mapping run consisted of 136 functional images (duration of 204 s). Eight prescan images (duration of 12 s) were discarded. During scanning, participants were required to perform a fixation task in which they had to press a button each time the fixation point turned from green to red. The average (std. err) performance on this task was 90.9% (±6.8%).

### MRI scanning and fMRI data processing

Scanning was carried out on a 3 Tesla Siemens Prisma MR-scanner using an 8-channel receiving SENSE head coil. A T1-weighted scan covering the whole-brain was recorded to chart each participant’s cortical anatomy. The functional scans were collected using standard EPI sequence (TR: 1500 ms; TE: 30 ms; voxel size of 3 mm isotropic, flip angle of 80 and a matrix size of 84 84 84 × 24). The T1-weighted whole-brain anatomical images were re-sampled to a 1 mm^3^ resolution. The resulting anatomical image was automatically segmented using Freesurfer (Dale et al., 1999) and subsequently edited manually. The cortical surface was reconstructed at the gray/white matter boundary and rendered on an inflated and smoothed 3D mesh (51).

The functional scans were analysed in the mrVista software package for MATLAB (available at http://white.stanford.edu/software). Head movement artifacts between and within functional scans were corrected (52). The functional scans were then averaged and coregistered to the anatomical scan (52) and interpolated to the anatomical segmentation. For comparison, the data was also analysed with conventional pRF modeling (Dumoulin and Wandell 2008). A 2D-gaussian model was fitted with parameters x_0_, y_0_, and σ where x_0_ and y_0_ are the receptive field center coordinates and σ is the spread (width) of the Gaussian signal, which is also the pRF size. We used SPM’s canonical difference of gammas for the HRF model. All parameter units are in degrees of visual angle and stimulus-referred. The borders of visual areas were determined on the basis of phase reversal (phase as obtained with the conventional pRF model). For each observer, six visual areas (V1, V2, V3, V4, LO1 and LO2) were manually delineated on the inflated cortical surface.

### Participants with albinism

Empirical data for this part of our study had previously been acquired as part of another study (24). In brief, a total of six patients with albinism and five aged-matched controls were included. Controls had normal or corrected-to-normal vision. The stimulus, procedure, data acquisition and preprocessing were identical to those reported by (23, 24). Retinotopic mapping was performed using three different stimuli: full field (FF); left field (LF) and right field (RF) stimulation. The study was performed monocularly: only the left eye was stimulated.

Analysis was performed using both conventional pRF modelling and our new MP approach. In the case of conventional pRF modelling, three models were used: a standard single Gaussian model and two bilateral Gaussian models, the latter two with positions that were symmetric in either the vertical or the horizontal axis. Three visual areas (V1 and V2 and V3) were defined in the left and right hemisphere of each observer. In observers A01 and A02, we could define V1 only in the right hemisphere due to too much noise in the phase maps.

The data of the healthy and albinism observers is available at XNAT central under the project ID: fMRI_micro_probing.

### Symmetry analysis of probe maps

For analysing the data of the observers with albinism, an additional symmetry analysis was developed based on the probe maps to quantify the degree of symmetry in the muRFs estimated for a voxel. This provides an indication of the degree to which visual information is misrouted. This analysis comprised three steps: 1) convert the scatter plot (figure 2A) into a heat map with a resolution of 40 × 40 bins (Figure 3B); 2) flip this “image” across the eight axes from 0 to 180 degrees in steps of 22.5 degrees, and 3) compute the correlation coefficient between the original and transformed (flipped) images. This resulted in correlation coefficients that indicate the extent to which the images are completely symmetrical (0) or identical (1).

Figure 7 depicts the symmetry estimation procedure for a typical voxel of an albinism observer. The degree of similarity between the original and mirrored image was translated into a correlation coefficient. By computing the symmetry coefficient in each probe map over the early visual cortex we identified regions that received input from both the contralateral and ipsilateral visual field.

**Figure 7.**
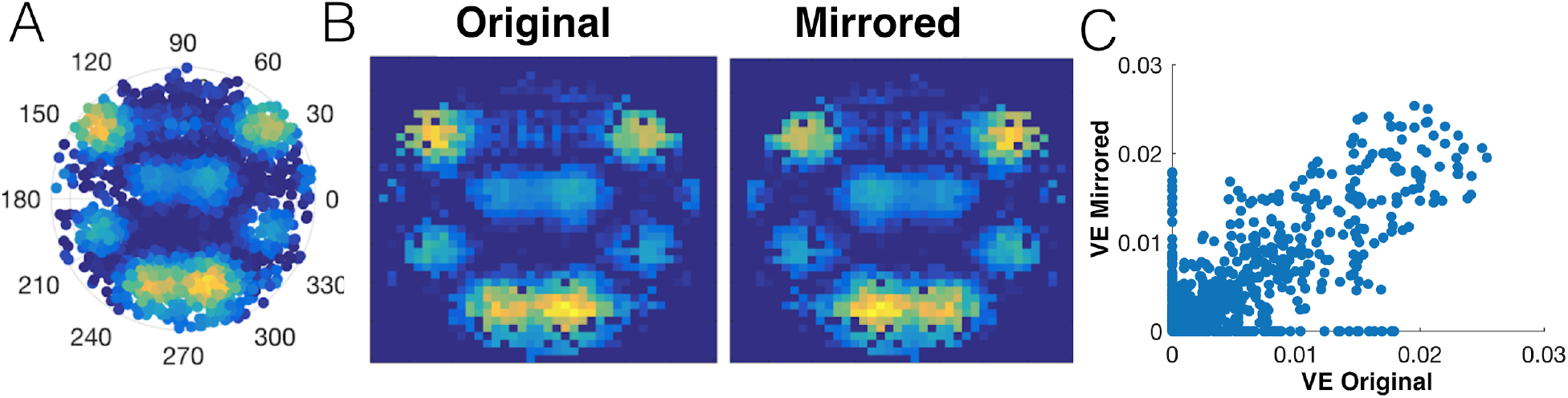
Calculation of a symmetry coefficient based on a probe map. A: Probe map of a typical V1-voxel of observer with albinism A01. B: Original and mirror images to the vertical meridian reconstructed. C: VE per bin of the mirrored image as function of the original image, for one representative voxel in V1. The symmetry coefficient of this particular voxel is 0.8.

## Acknowledgments

We thank Koen V. Haak for suggesting to apply MP to albinism data. Authors JC, KA and AI were supported by the European Union’s Horizon 2020 research and innovation programme under the Marie Sklodowska-Curie grant agreements No. 641805 (NextGenVis) and No. 661883 (EGRET) awarded to FWC and MBH. The funding organization had no role in the design, conduct, analysis, or publication of this research.

## Supplementary Material

### Simulations

To verify the accuracy of our model, we simulated multiple pRFs within a voxel using the conventional pRF model. These were centered at multiple locations and had different sizes, based on equation 1. The total profile was given by the sum of the individuals muRFs simulated. Next, the simulated time series were calculated based on the steps to generate the predicted times series, equation 9. We used the standard moving bar stimuli, described in the stimulus section. White gaussian noise was added to the simulated time series.

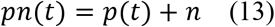

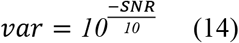

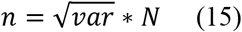

Where *pn*(*t*) is the predicted time series with added noise, *p*(*t*) is the predicted time series and *n* is the white gaussian noise, while *var* is the variance of the noise.

The first simulation was aimed at evaluating the behaviour of MP and its robustness to noise. To do so, per simulation we defined two muRFs (1 deg size) mirrored across different meridians. To each simulation, white gaussian noise was added to the simulated time series, such that the SNR levels varied between 0.2 and 10. A total 144 simulations (24 different position × 6 levels of noise) were generated. The second set of simulations was aimed at evaluating how the number of probes retained affects the accuracy in position and size estimation of the muRFs. Per simulation, we varied the k-threshold (percentage of micro probes retained) from 1% (more restrictive) to 100 % in steps of 1%. We simulated one pRF in with six different pRF sizes, across 30 different positions. The third set of simulations was aimed at evaluating the method’s ability to recover the proper number of muRFs. Per simulation, the number of muRFs could assume any random integer value between one and four. Each of these muRFs could be randomly positioned within the visual field. A total of 1000 simulations were performed using a SNR of 0.5 and a muRF size of 1.

### Effect of k-threshold

Figure S1 shows the eccentricity, polar angle and size error as function of the k-threshold. In general restrictive k-thresholds minimize the eccentricity error, while more lenient ones minimize the pRF size error. The polar angle estimation is not influenced by the k-threshold. Figure S1 shows the eccentricity, polar angle and size error for the six different sizes separately.

**Figure S1.**
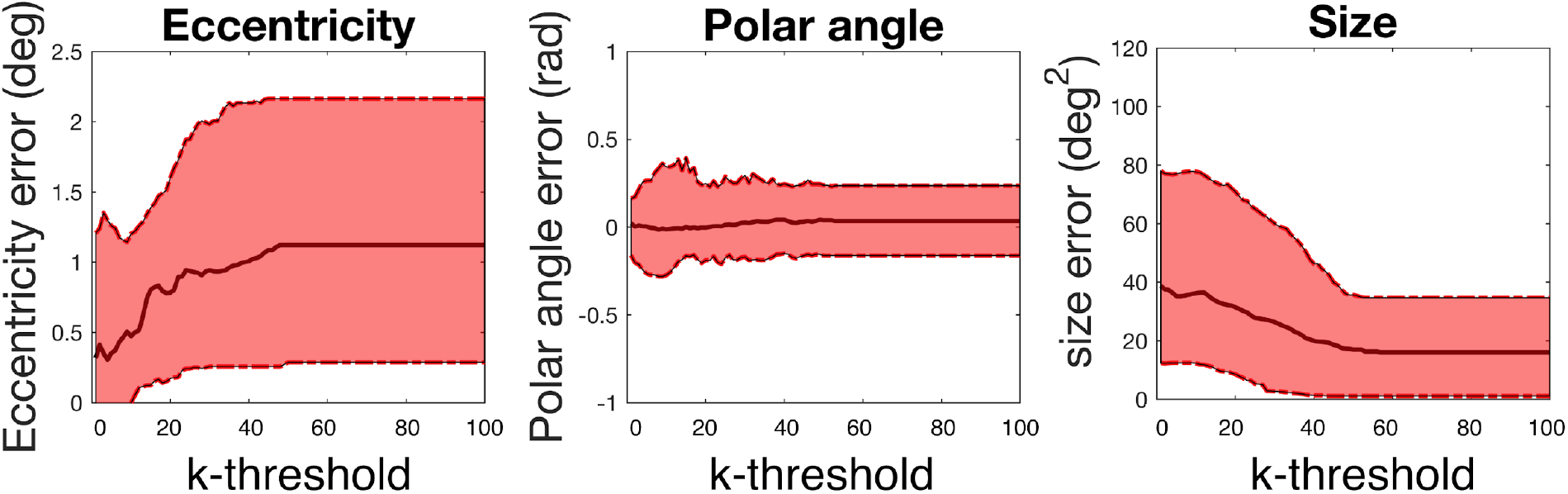
Effect of the k-threshold in the estimated muRFs properties. Eccentricity, polar angle and size error between simulated and estimated pRFs as a function of the variance explained cutoff. The black continuous line represents the median Error bars represent the 5% and 95 % confidence interval.

### Robustness to noise

Figure S2 (panel A) shows the result of MP for a single simulated bilateral pRF mirrored to the vertical meridian. The probes were positioned at [3,3; 3,−3]. Note that probes with the simulated muRFs present a higher variance explained than the remaining visual field. After thresholding the probe maps, the two simulated clusters were recovered (panel B).

Figure S2 (panel C) compares time series as predicted by MP (red) and conventional pRF (blue). At all noise levels, MP well-captured the dynamics of the simulated times series, and fitted the simulated data better than the conventional pRF (panel D). MP accurately detects the position of the muRFs however it underestimates its size (panel E and F). While, in the majority of cases the conventional pRF accurately detects only one of the muRFs, in some cases, the conventional pRF model increases its size in an attempt to improve the fitting (panel F). The accuracy of MP to recover the position and size of the muRFs is independent of the noise level (panel E and F).

**Figure S2.**
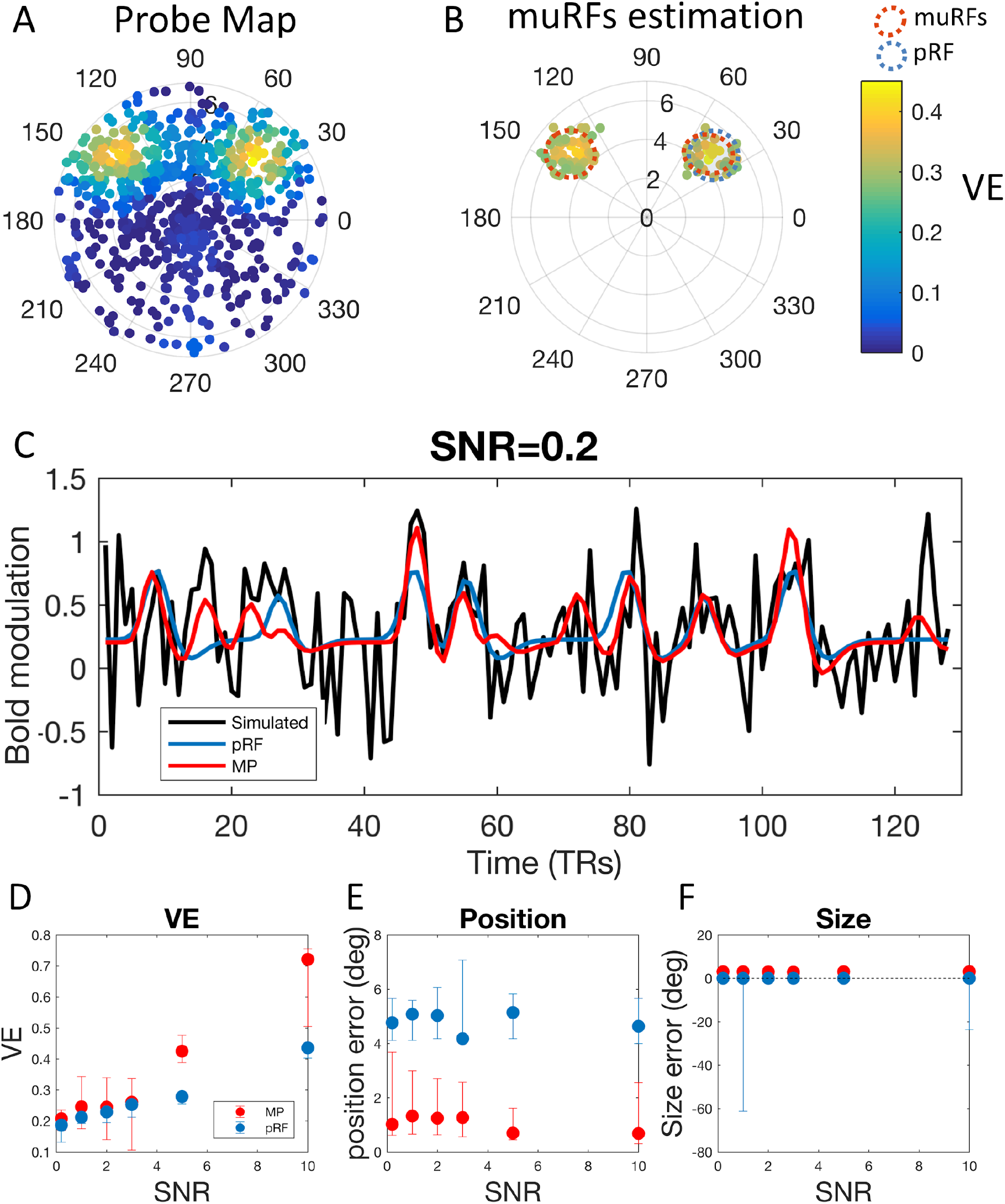
Microprobing simulated bilateral pRFs. Panel A: probe map obtained from the simulated time series. The colour bar represent the variance explained. Panel B: Thresholded probe map, using a k threshold of 15% and the results of the clustering and gaussian mixture model analysis. The estimated muRFs and pRF are indicated by the shaded gray gaussians, outlined in red and blue respectively. Panel C: simulated time series of bilateral pRFs (black) and the predicted time series estimated with conventional pRF (blue) and with MP (red), for the six levels of SNR tested. The predicted time series estimated on the basis of MP was calculated using the estimated muRF. Panels D, E and F represent the VE, pRF/muRF positional error and pRF/muRF size error as function of the SNR, respectively. The muRFs estimated position and size closely resemble what was simulated (median VE, positional and size error and VE aremuRF:0.69, 0.4 deg and 2.95 deg2; pRF:0.42, 4.63 deg and 0 deg2, respectively at a SNR of 10). The mean position error calculated as the euclidean distance between the simulated and estimated muRFs. Even at very low SNR (0.2), MP could accurately detect the position of the muRF. The size error corresponds to the mean difference in area between the simulated and estimated muRFs.

### Factors affecting MP performance

We find that the main factors affecting the accuracy of MP are the number of muRFs and their distance. Figure S3 shows that the positional error of the estimated muRF is approximately constant over eccentricity. However, this positional error does depend on the number of muRFs. Figure S3 shows that while for one muRF the method is accurate, for multiple muRFs the positional error increases. The method’s ability to determine the actual number of muRFs improves with the distance between simulated muRFs (figure S3).

The main factors affecting the MP performance are the number of muRFs and distance between the muRF. When the muRFs are too close and their activity superimposes, the muRFs individual characteristics cannot be captured.

**Figure S3.**
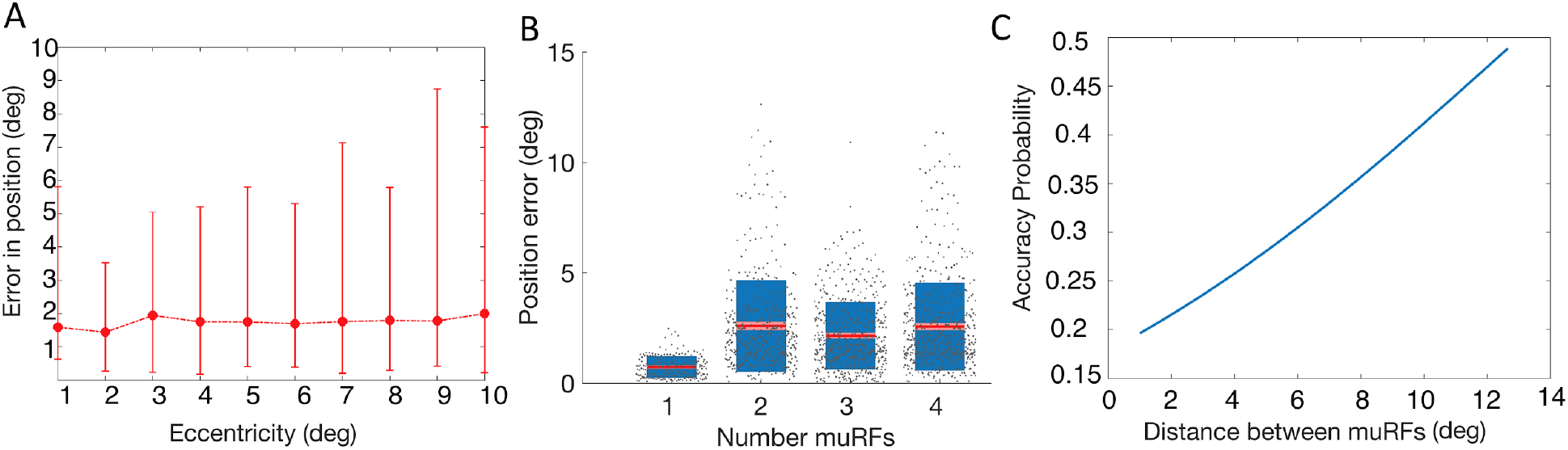
A: Positional error (in euclidean distance) between simulated and estimated muRF as a function of eccentricity. Error bars represent the 5% and 95 % confidence interval. B: Box plot of the positional error as a function of the number of simulated muRFs. C: Probability density function of accurately detecting all the simulated muRFs as a function of their distance (deg).

### Supplementary data on human observers

Figure S4 represents muRF eccentricity as function of pRF eccentricity in human observers. The muRFs detected within a voxel were sorted from the most foveal (represented in blue) to the most peripheral (purple). While in the fovea the muRFs eccentricities vary substantially, in the periphery they largely overlap. For voxels with multiple muRFs, in the fovea (<2 degrees) the muRFs eccentricity estimates are larger, i.e. more peripheral, than those of the pRF. This can be related to the finding that the muRFs in the fovea are more elongated and with statistical biases. Note that not all voxels contained the same number of muRFs-very few voxels got assigned four muRFs, hence the data for these is much noisier and trends are less clear.

**Figure S4:**
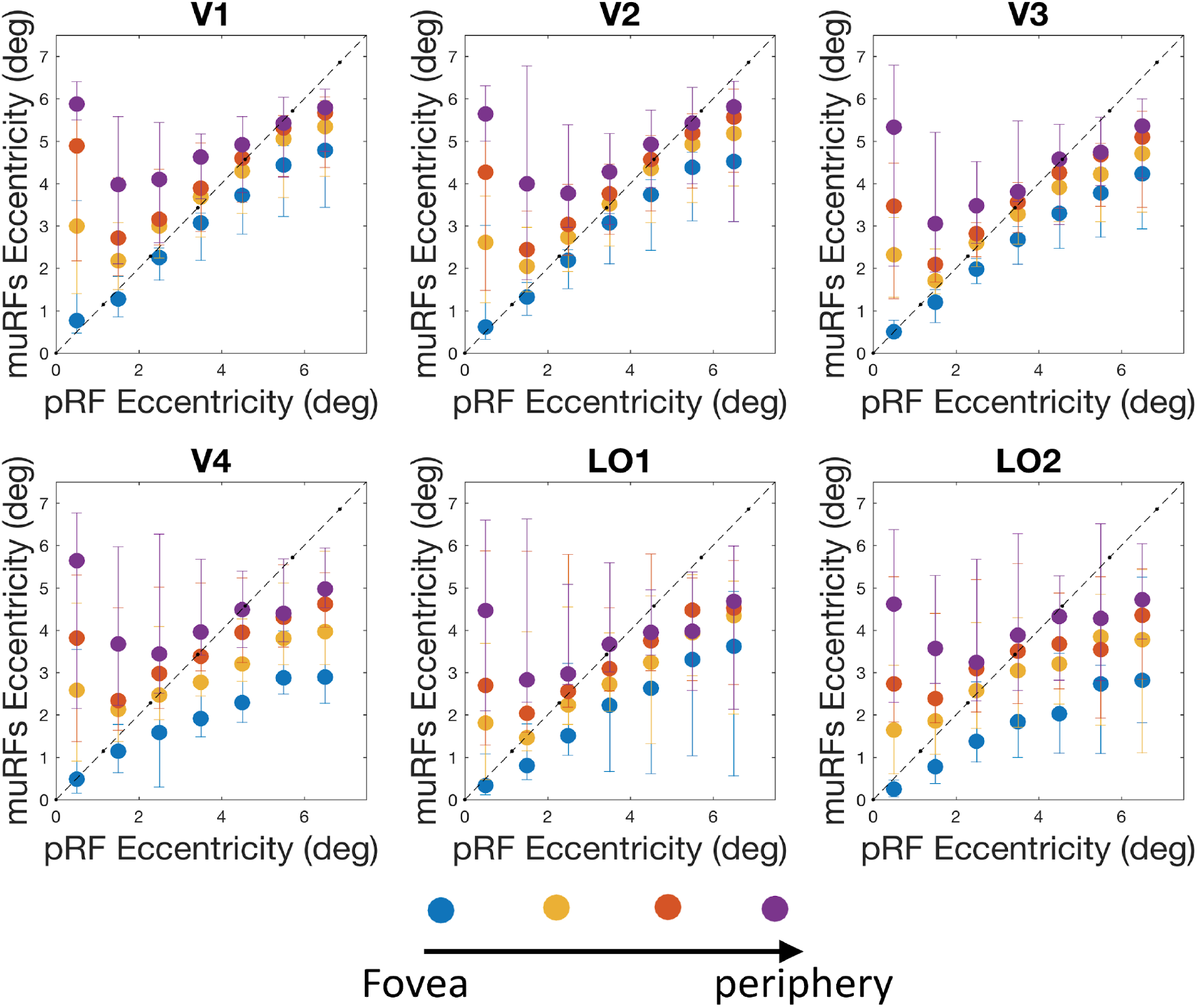
Median eccentricity of the muRFs as a function of the eccentricity of the conventional pRF (the dashed line represents a perfect correlation). The muRFs eccentricities were sorted per voxel from the most foveal to the most peripheral muRF. The muRFs eccentricities were then binned in 1 deg bins of eccentricity, representing the data of 7 healthy observers (14 hemispheres). Error bars represents 5% and 95% confidence interval.

**Table. S1.** Represent the correlation factor and the corresponding significance value for the most foveal to the most peripheric muRF, for all the visual areas analysed.

| Correlation Coefficient |  |  |  |  | p-value |  |  |  |
| --- | --- | --- | --- | --- | --- | --- | --- | --- |
|  | 1st muRF | 2nd muRF | 3rd muRF | 4th muRF | 1st muRF | 2nd muRF | 3rd muRF | 4th muRF |
| V1 | 0.99 | 0.92 | 0.79 | 0.56 | 0.0000 | 0.0036 | 0.0341 | 0.1951 |
| V2 | 0.99 | 0.95 | 0.87 | 0.56 | 0.0000 | 0.0013 | 0.0109 | 0.1897 |
| V3 | 0.99 | 0.96 | 0.90 | 0.71 | 0.0000 | 0.0005 | 0.0061 | 0.0748 |
| V4 | 1.00 | 0.92 | 0.84 | 0.46 | 0.0000 | 0.0030 | 0.0168 | 0.2949 |
| LO1 | 1.00 | 0.98 | 0.94 | 0.77 | 0.0000 | 0.0001 | 0.0015 | 0.0434 |
| LO2 | 0.98 | 0.96 | 0.96 | 0.56 | 0.0001 | 0.0008 | 0.0006 | 0.1952 |

Figure S5 shows the muRF size as function of its eccentricity. The muRFs were sorted from the smallest (blue) to the largest (purple). Although with shallow slopes the muRF size increases with eccentricity for all the visual areas. This slope becomes more steep as we go through the cortical hierarchy.

**Figure S5:**
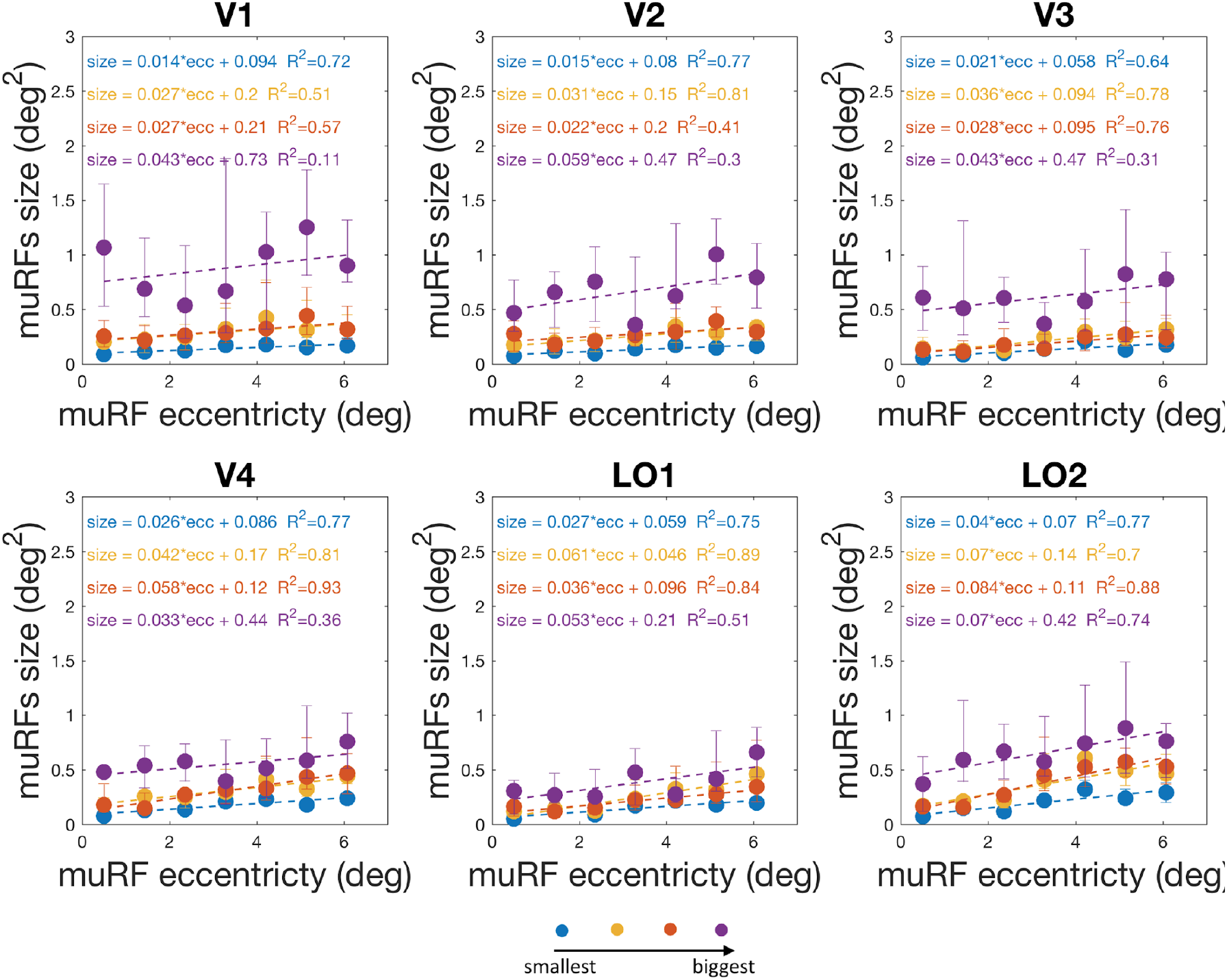
Median muRFs size as a function of eccentricity of the conventional pRF. The muRF size was calculated using a k-threshold of 15% and VEr of 0.1. The muRF size was binned in 1 deg bins of eccentricity, and it represents the data of 7 healthy observers (14 hemispheres). Error bars represents 25% and 75% confidence interval. The dashed lines represent the linear fit and the corresponding equations and goodness of fit are shown on top of the graph.

Figure S6 shows two probe maps where pRF estimated resulted in a higher VE than muRFs. It is noticeable that the pRFs are larger than the muRFs. Panel B shows that the increase in size results in a higher variance explained.

**Figure S6.**
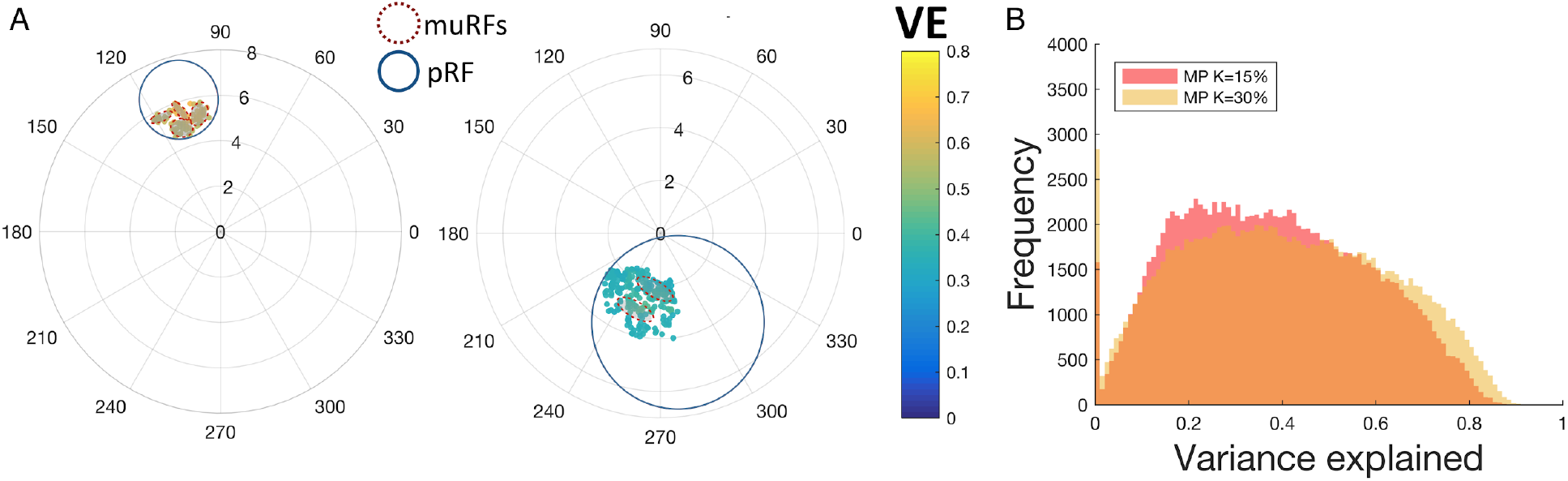
Panel A: probe maps of V1 voxels extracted from the right hemisphere of observer S07 (k-threshold: 15%, VEr: 0.1). The muRFs and pRFs estimated are outlined with a dashed red and continuous blue line respectively. Data obtained during retinotopic mapping. Panel B: Histogram of the VE for muRFs obtained using a k-threshold of 15% (red) and 30% (yellow). The VE is based on the cumulative activity of the number of muRFs. Each histogram represents the data of the six visual areas, of 7 healthy observers(14 hemispheres), distributed over 100 bins.

**Figure S7.**
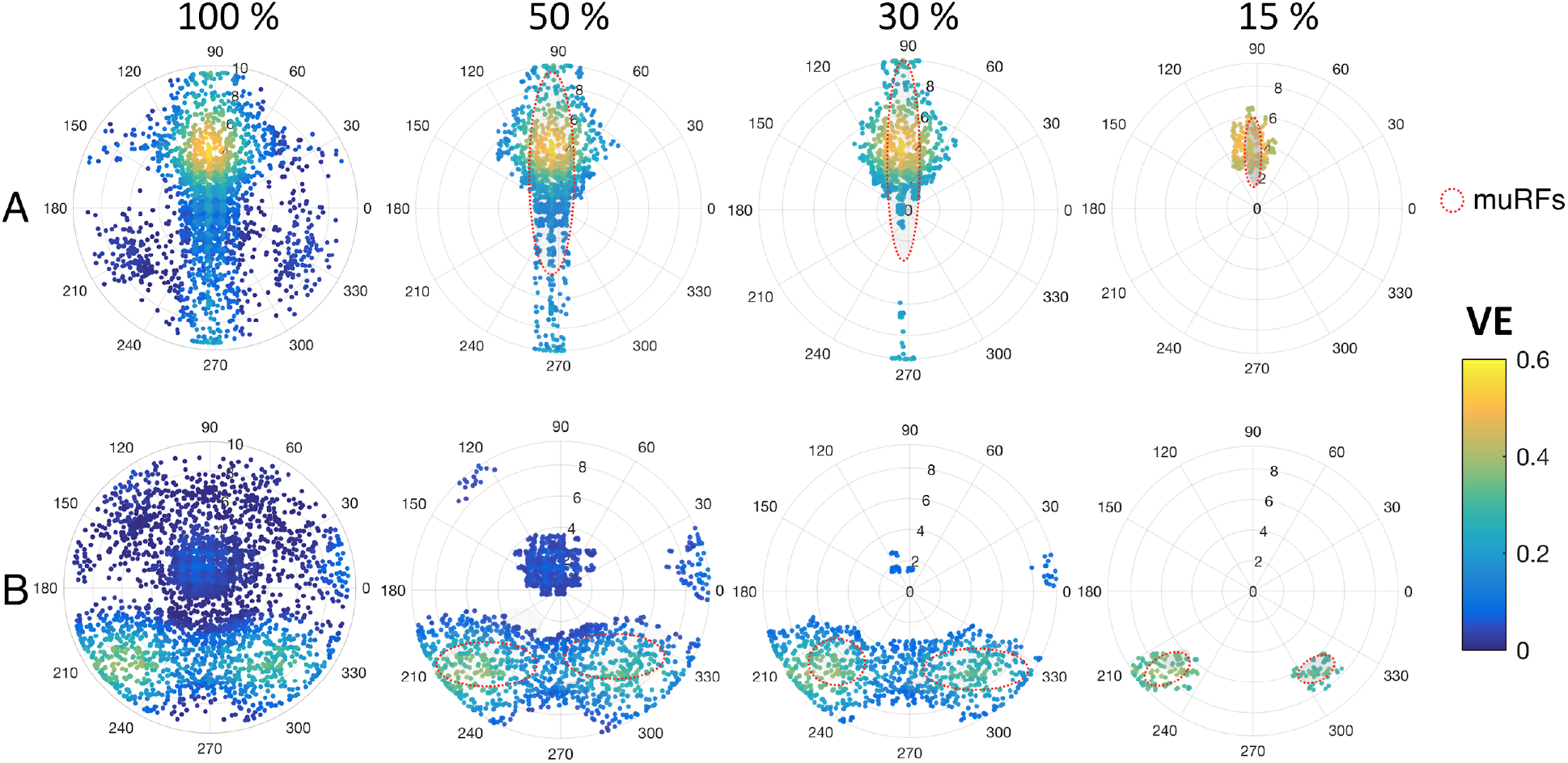
Two V1 voxels probe maps thresholded at 100%, 50%, 30% and 15%, the MP-based muRF estimates have dashed red outlines. Panel A corresponds to a healthy control and panel B to an observer with albinism.

**Figure S8.**
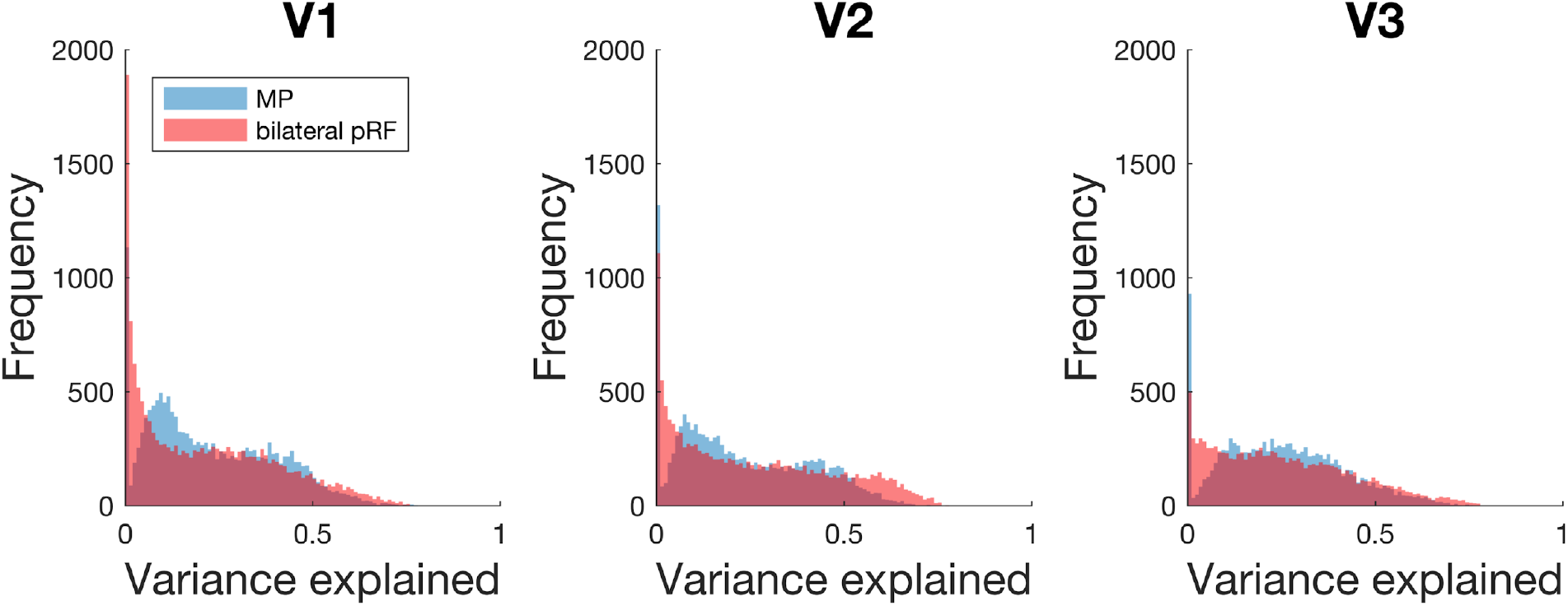
Histogram of the VE for muRFs and pRFs for several visual areas, represented with blue and red respectively. The VE based on the cumulative activity of the number of muRFs. Each histogram represents the data of 7 healthy observers (14 hemispheres).

**Figure S9.**
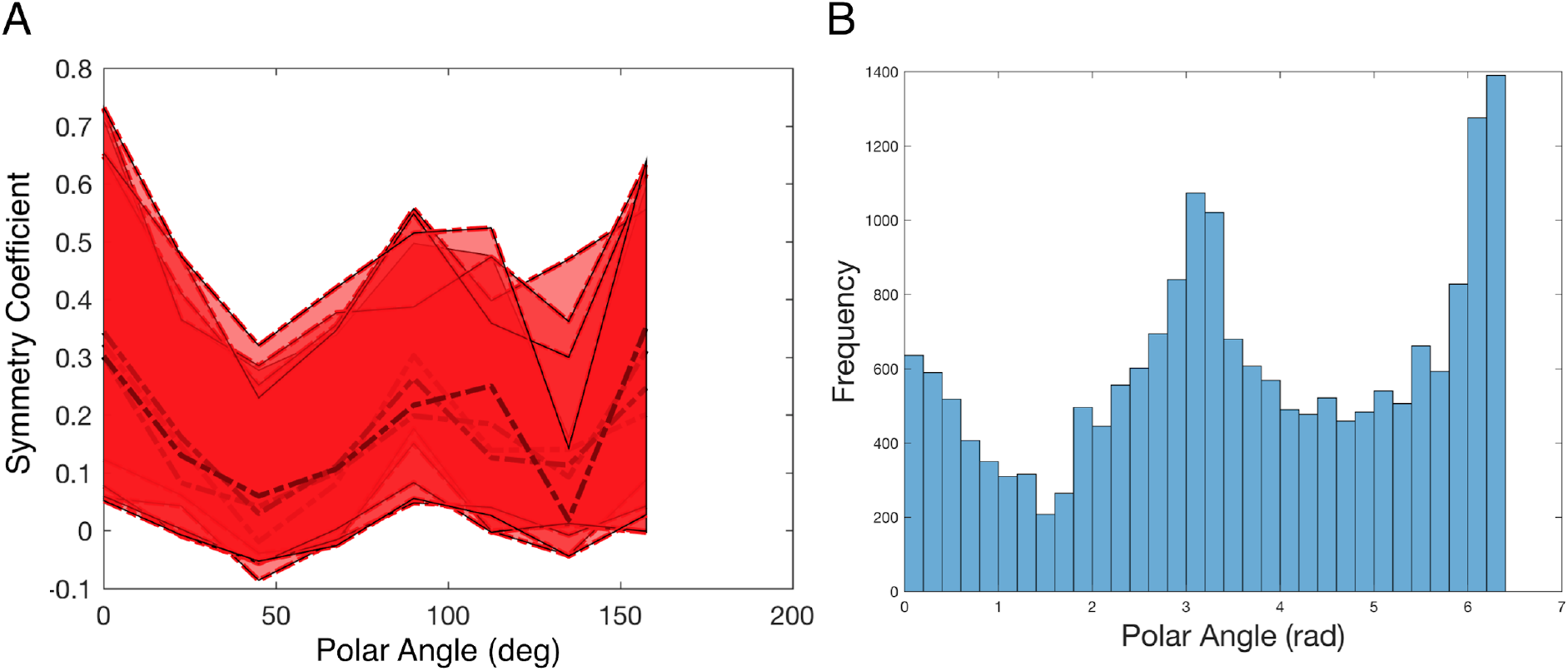
Panel A Symmetry coefficients for every healthy control observer plotted individually. Symmetry coefficients for the V1 of the right hemisphere during full field stimulation were calculated across 8 symmetry axes, from 0 to *180* deg in steps of 22.5 deg. The dashed lines represent the the 5%, 50% and 95% confidence intervals, for every participant. Panel B: Histogram of the estimated polar angle of voxels obtained with the conventional pRF method. The histogram represent the data for three visual areas (V1, V2, V3), of five controls (10 hemispheres).

**Table. S2.** Misrouting of albinism and controls. The mean misrouting extent (deg) previously determined by (24), and misrouting coefficient calculated with MP-average symmetry coefficient to 90 deg across the entire hemisphere - based on the full field condition. C refers to the median of the controls coefficients.

| Observer | Mean misrouting extent (deg) | Misrouting coefficient |  |  |
| --- | --- | --- | --- | --- |
|  |  | V1 | V2 | V3 |
| A01 | >9.0 | 0.57 | 0.66 | 0.69 |
| A02 | >9.0 | 0.72 | 0.73 | 0.73 |
| A03 | >9.0 | 0.65 | 0.72 | 0.77 |
| A04 | 5.2 | 0.2 | 0.29 | 0.29 |
| A05 | 4.2 | 0.16 | 0.03 | 0.09 |
| A06 | 3.1 | 0.34 | 0.48 | 0.44 |
| C | 0 | 0.27 | 0.27 | 0.21 |

Figure S10A represents a projection of the number of muRFs on the visual cortex. In both controls and Albinism the number of muRFs increases with eccentricity, this effect if present in the three conditions and visual areas tested S10B. Moreover the number of muRFs is significantly higher for Albinism when compared to age matched controls for the full field condition, in the three visual areas tested. For LF and RF there is no difference in the number of muRFs between the two population groups. As expected the maps of the number muRFs follow the same spatial organization as the symmetry analysis.

**Figure S10.**
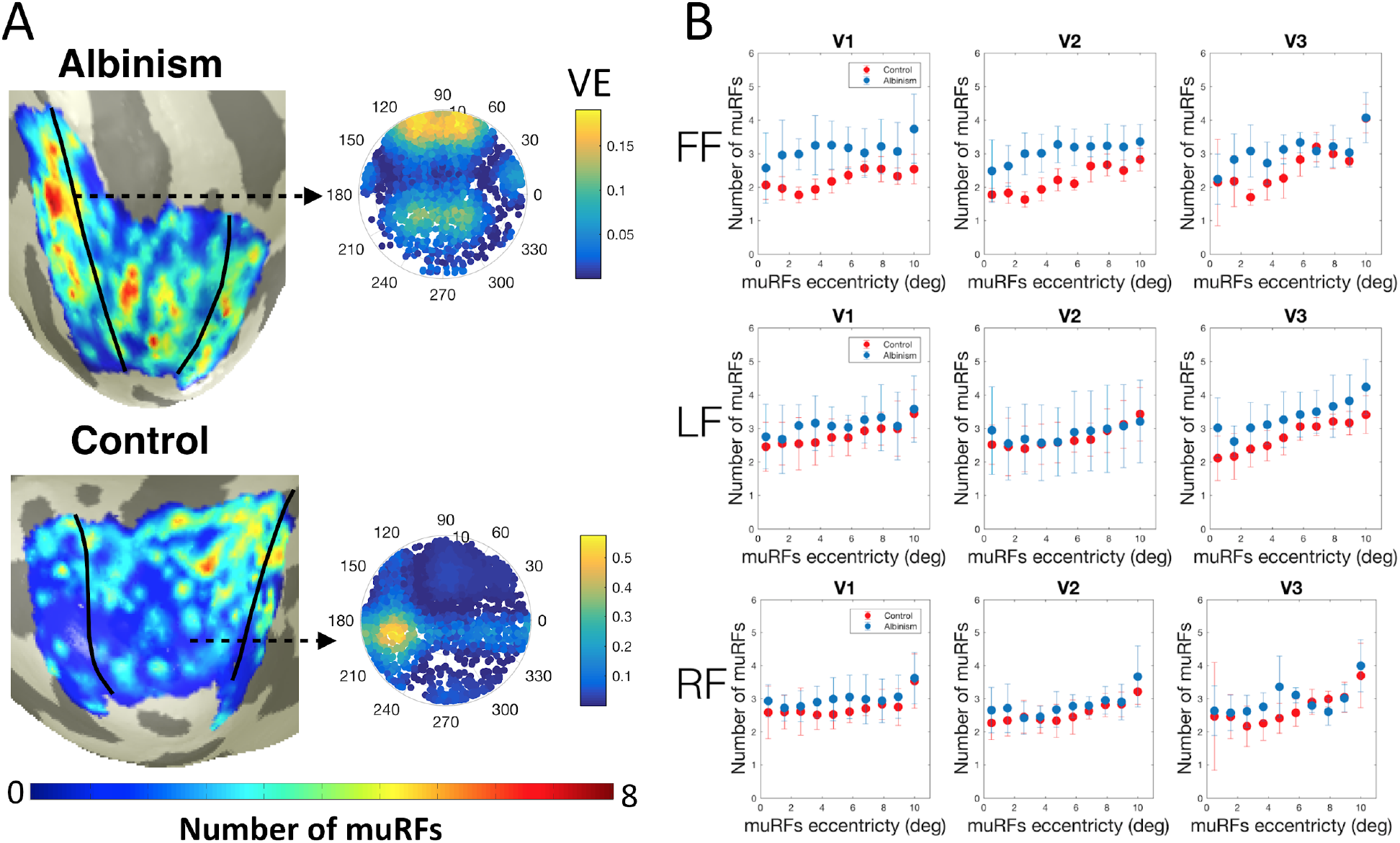
A Projection on an inflated brain mesh of the number of muRFs, right hemispheres of the observer with albinism A03 and S07. B: Median of the number of muRFs binned in 1 degree bins as function of the muRFs eccentricity estimated, for the 3 visual areas and the 3 stimuli conditions tested. Albinism and aged matched controls are represented in red and blue respectively, the error bars represent the 25% and 75 % confidence interval.

**Figure S11.**
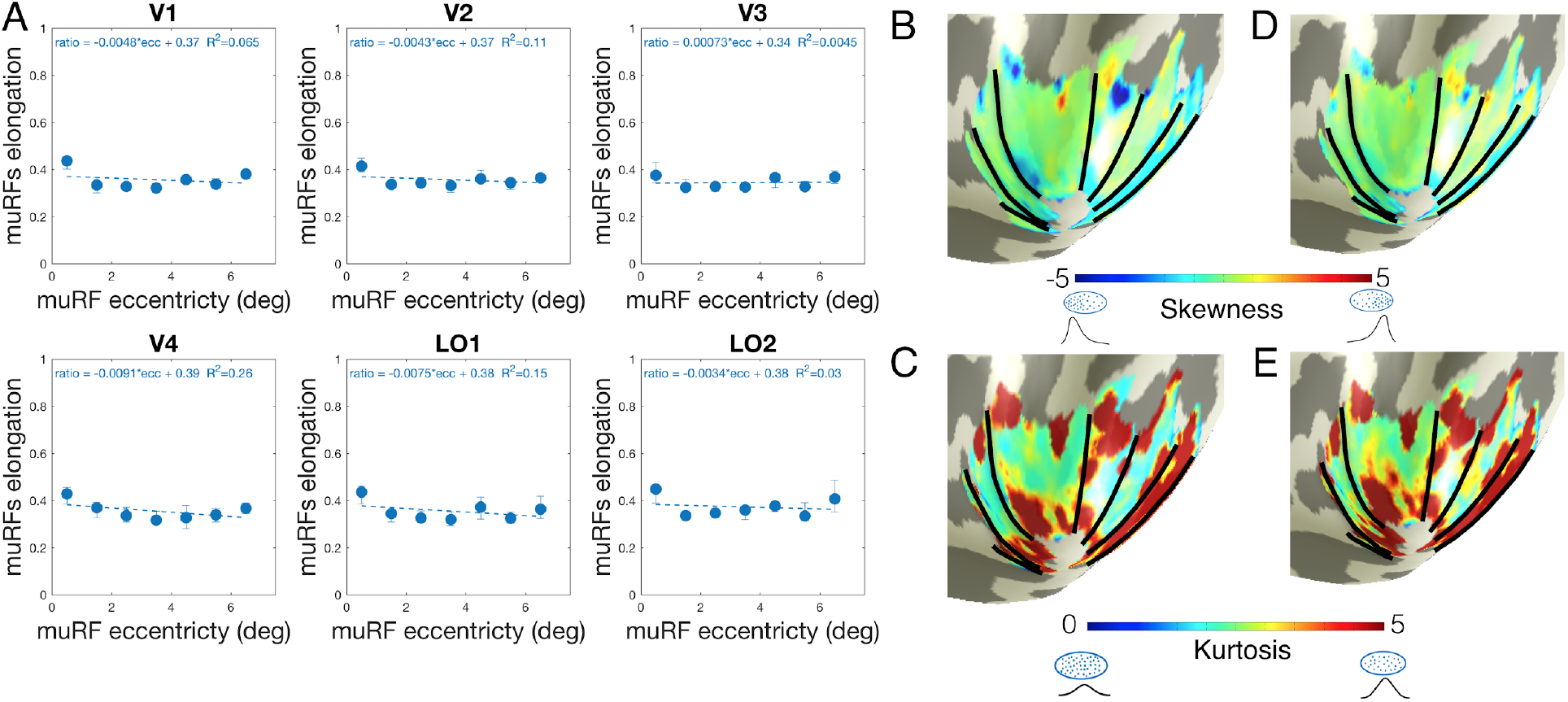
Shape characterization of muRFs. A Median of the muRF elongation size as function of eccentricity. The muRF elongation was calculated as one minus the ratio between the minor and major axis of the muRFs. Error bars represent the 5% and 95% confidence interval.The dashed line represents the linear fit. represent the data of 14 hemispheres of 7 observers. Projection on an inflated brain mesh of the skewness (B) and kurtosis of muRFs (right hemisphere of participant S07). Skewness was calculated across the main axis of one (arbitrarily) chosen muRF. Black lines indicate the borders of visual areas. D and E show the skewness and kurtosis across the secondary axis of one (arbitrarily) chosen muRF.

## References

1. Hubel DH, Wiesel TN (1974) Uniformity of monkey striate cortex: a parallel relationship between field size, scatter, and magnification factor. J Comp Neurol 158(3):295–305.

2. Hubel DH, Wiesel TN (1974) Sequence regularity and geometry of orientation columns in the monkey striate cortex. J Comp Neurol 158(3):267–293.

3. Hubel DH, Wiesel TN, Stryker MP (1977) Orientation columns in macaque monkey visual cortex demonstrated by the 2-deoxyglucose autoradiographic technique. Nature 269(5626):328–330.

4. Dumoulin SO, Wandell BA (2008) Population receptive field estimates in human visual cortex. Neuroimage 39(2):647–660.

5. Wandell BA, Smirnakis SM (2009) Plasticity and stability of visual field maps in adult primary visual cortex. Nat Rev Neurosci 10(12):873–884.

6. Haak KV, Cornelissen FW, Morland AB (2012) Population receptive field dynamics in human visual cortex. PLoS One 7(5):e37686.

7. Dumoulin SO, Knapen T (2018) How Visual Cortical Organization Is Altered by Ophthalmologic and Neurologic Disorders. Annu Rev Vis Sci 4:357–379.

8. Haak KV, et al. (2013) Connective field modeling. Neuroimage 66:376–384.

9. Harvey BM, Fracasso A, Petridou N, Dumoulin SO (2015) Topographic representations of object size and relationships with numerosity reveal generalized quantity processing in human parietal cortex. Proc Natl Acad Sci U S A 112(44):13525–13530.

10. Thomas JM, et al. (2015) Population receptive field estimates of human auditory cortex. Neuroimage 105:428–439.

11. Finlay BL, Schiller PH, Volman SF (1976) Meridional differences in orientation sensitivity in monkey striate cortex. Brain Res 105(2):350–352.

12. DeValois RL (1982) Early Visual Processing: Feature Detection or Spatial Filtering? Lecture Notes in Biomathematics, pp 152–174.

13. Ringach DL (2002) Spatial structure and symmetry of simple-cell receptive fields in macaque primary visual cortex. J Neurophysiol 88(1):455–463.

14. Chapin JK (1986) Laminar differences in sizes, shapes, and response profiles of cutaneous receptive fields in the rat SI cortex. Exp Brain Res 62(3):549–559.

15. Merkel C, Hopf J-M, Schoenfeld MA (2018) Spatial elongation of population receptive field profiles revealed by model-free fMRI back-projection. Hum Brain Mapp 39(6):2472–2481.

16. Silson EH, Reynolds RC, Kravitz DJ, Baker CI (2018) Differential Sampling of Visual Space in Ventral and Dorsal Early Visual Cortex. J Neurosci 38(9):2294–2303.

17. Keliris GA, Li Q, Papanikolaou A, Logothetis NK, Smirnakis SM (2019) Estimating average single-neuron visual receptive field sizes by fMRI. Proc Natl Acad Sci U S A 116(13):6425–6434.

18. Baseler HA, et al. (2011) Large-scale remapping of visual cortex is absent in adult humans with macular degeneration. Nat Neurosci 14(5):649–655.

19. Papanikolaou A, et al. (2014) Population receptive field analysis of the primary visual cortex complements perimetry in patients with homonymous visual field defects. Proc Natl Acad Sci U S A 111(16):E1656–65.

20. Hoffmann MB, Dumoulin SO (2015) Congenital visual pathway abnormalities: a window onto cortical stability and plasticity. Trends Neurosci 38(1):55–65.

21. Hoffmann MB, Tolhurst DJ, Moore AT, Morland AB (2003) Organization of the visual cortex in human albinism. J Neurosci 23(26):8921–8930.

22. Kaule FR, et al. (2014) Impact of chiasma opticum malformations on the organization of the human ventral visual cortex. Hum Brain Mapp 35(10):5093–5105.

23. Hoffmann MB, et al. (2012) Plasticity and stability of the visual system in human achiasma. Neuron 75(3):393–401.

24. Ahmadi K, et al. Population receptive field and connectivity properties of the early visual cortex in human albinism. doi:10.1101/627265.

25. Hubel DH, Wiesel TN (1967) Cortical and callosal connections concerned with the vertical meridian of visual fields in the cat. J Neurophysiol 30(6):1561–1573.

26. Makarov VA, Schmidt KE, Castellanos NP, Lopez-Aguado L, Innocenti GM (2008) Stimulus-dependent interaction between the visual areas 17 and 18 of the 2 hemispheres of the ferret (Mustela putorius). Cereb Cortex 18(8):1951–1960.

27. Schmidt KE, Lomber SG, Innocenti GM (2010) Specificity of neuronal responses in primary visual cortex is modulated by interhemispheric corticocortical input. Cereb Cortex 20(12):2776–2786.

28. Choudhury BP, Whitteridge D, Wilson ME (1965) The function of the callosal connections of the visual cortex. Q J Exp Physiol Cogn Med Sci 50:214–219.

29. Raiguel S, et al. (1997) Size and shape of receptive fields in the medial superior temporal area (MST) of the macaque. Neuroreport 8(12):2803–2808.

30. Ungerleider LG, Desimone R (1986) Projections to the superior temporal sulcus from the central and peripheral field representations of V1 and V2. J Comp Neurol 248(2):147–163.

31. Saito H, et al. (1986) Integration of direction signals of image motion in the superior temporal sulcus of the macaque monkey. J Neurosci 6(1):145–157.

32. Yildirim F, Carvalho J, Cornelissen FW (2018) A second-order orientation-contrast stimulus for population-receptive-field-based retinotopic mapping. Neuroimage 164:183–193.

33. Greene CA, Dumoulin SO, Harvey BM, Ress D (2014) Measurement of population receptive fields in human early visual cortex using back-projection tomography. J Vis 14(1). doi:10.1167/14.1.17.

34. Dukart J, Bertolino A (2014) When Structure Affects Function – The Need for Partial Volume Effect Correction in Functional and Resting State Magnetic Resonance Imaging Studies. PLoS ONE 9(12):e114227.

35. Zhou Y, Thompson PM, Toga AW (1999) Extracting and Representing the Cortical Sulci. IEEE Comput Graph Appl 19(3):49–55.

36. Fracasso A, et al. (2016) Lines of Baillarger in vivo and ex vivo: Myelin contrast across lamina at 7 T MRI and histology. NeuroImage 133:163–175.

37. Le R, Witthoft N, Ben-Shachar M, Wandell B (2017) The field of view available to the ventral occipito-temporal reading circuitry. J Vis 17(4):6.

38. Klein BP, Harvey BM, Dumoulin SO (2014) Attraction of position preference by spatial attention throughout human visual cortex. Neuron 84(1):227–237.

39. Binda P, Thomas JM, Boynton GM, Fine I (2013) Minimizing biases in estimating the reorganization of human visual areas with BOLD retinotopic mapping. J Vis 13(7):13.

40. Senden M, Reithler J, Gijsen S, Goebel R (2014) Evaluating population receptive field estimation frameworks in terms of robustness and reproducibility. PLoS One 9(12):e114054.

41. Zuiderbaan W, Harvey BM, Dumoulin SO (2012) Modeling center-surround configurations in population receptive fields using fMRI. J Vis 12(3):10.

42. Fracasso A, Luijten PR, Dumoulin SO, Petridou N (2018) Laminar imaging of positive and negative BOLD in human visual cortex at 7 T. Neuroimage 164:100–111.

43. Shmuel A, Yacoub E, Chaimow D, Logothetis NK, Ugurbil K (2007) Spatio-temporal point-spread function of fMRI signal in human gray matter at 7 Tesla. Neuroimage 35(2):539–552.

44. Yacoub E, Harel N, Ugurbil K (2008) High-field fMRI unveils orientation columns in humans. Proc Natl Acad Sci U S A 105(30):10607–10612.

45. Adaszewski S, Slater D, Melie-Garcia L, Draganski B, Bogorodzki P (2018) Simultaneous estimation of population receptive field and hemodynamic parameters from single point BOLD responses using Metropolis-Hastings sampling. Neuroimage 172:175–193.

46. Zeidman P, Silson EH, Schwarzkopf DS, Baker CI, Penny W (2018) Bayesian population receptive field modelling. Neuroimage 180(Pt A):173–187.

47. Azzopardi P, Cowey A (1993) Preferential representation of the fovea in the primary visual cortex. Nature 361(6414):719–721.

48. Boynton GM, Demb JB, Heeger DJ (1996) fMRI responses in human V1 correlate with perceived stimulus contrast. Neuroimage 3(3):S265.

49. Friston KJ (1998) Modes or models: a critique on independent component analysis for fMRI. Trends Cogn Sci 2(10):373–375.

50. Davies DL, Bouldin DW (1979) A cluster separation measure. IEEE Trans Pattern Anal Mach Intell 1(2):224–227.

51. Wandell BA, Chial S, Backus BT (2000) Visualization and measurement of the cortical surface. J Cogn Neurosci 12(5):739–752.

52. Nestares O, Heeger DJ (2000) Robust multiresolution alignment of MRI brain volumes. Magn Reson Med 43(5):705–715.

